# Inference of emergent spatio-temporal processes from single-cell sequencing reveals feedback between *de novo* DNA methylation and chromatin condensates

**DOI:** 10.1101/2020.12.30.424823

**Authors:** Fabrizio Olmeda, Tim Lohoff, Stephen J Clark, Laura Benson, Felix Krüger, Wolf Reik, Steffen Rulands

**Author notes:** Joint corresponding authors: **Corresponding authors email addresses:**.

## Abstract

Recent breakthroughs in single-cell genomics allow probing molecular states of cells with unprecedented detail along the sequence of the DNA. Biological function relies, however, on emergent processes in the three-dimensional space of the nucleus, such as droplet formation through phase separation. Here, we use single-cell multi-omics sequencing to develop a theoretical framework to rigorously map epigenome profiling along the DNA sequence onto a description of the emergent spatial dynamics in the nucleus. Drawing on scNMT-seq multi-omics sequencing *in vitro* and *in vivo* we exemplify our approach in the context of exit from pluripotency and global *de novo* methylation of the genome. We show how DNA methylation patterns of the embryonic genome are established through the interplay between spatially correlated DNA methylation and topological changes to the DNA. This feedback leads to the predicted formation of 30-40nm sized condensates of methylated DNA and determines genome-scale DNA methylation rates. We verify these findings with orthogonal single cell multi-omics data that combine the methylome with HiC measurements. Notably, this scale of chromatin organization has recently been described by super-resolution microscopy. Using this framework, we identify local methylation correlations in gene bodies that precede transcriptional changes at the exit from pluripotency. Our work provides a general framework of how mechanistic insights into emergent processes underlying cell fate decisions can be gained by the combination of single-cell multi-omics and methods from theoretical physics that have not been applied in the context of genomics before.

**Highlights:**

- We develop methodology to infer collective spatio-temporal processes in the physical space of the nucleus from single-cell methylome sequencing experiments.
- We show that DNA methylation relies on a feedback between *de novo* methylation and nanoscale changes in DNA topology, leading to the formation of methylation condensates.
- Chromatin condensates at this scale have recently been described by high-resolution microscopy but have remained without mechanistic explanation.
- Using this framework, we identify changes in the distribution of DNA methylation marks in gene bodies that precede gene silencing at the exit from pluripotency.

## Introduction

The organisation of cells into complex tissues relies on tightly regulated molecular and cellular programs. On the molecular level, different layers are involved in regulating cellular fate, such as transcription factor networks, dynamic changes in the topology of the DNA and chemical modifications of DNA and histones. Recent technological breakthroughs in single-cell genomics now allow probing these processes with unprecedented detail, giving access to detailed molecular profiles of single cells, such as the expression of thousands of genes, to epigenetic modifications of the DNA at single loci and to the spatial organisation of chromatin (*1*). Single-cell multi-omics technologies unlock molecular profiles of several layers of regulation in the same cell (*2*–*4*). These technological developments, along with computational methods, have led to detailed descriptions of the molecular states of cells (*5*). Biological function, however, is determined by emergent (collective) processes on the cellular and tissue scale, which cannot be straightforwardly inferred from the multitude of interacting processes on the molecular scale (*6*). Further, while sequencing typically gives information on linear measurements along the one-dimensional sequence of the DNA (sequence space), such collective processes often rely on the spatial arrangement and dynamics of molecules in the three-dimensional space of the nucleus (physical space) such as in phase separation phenomena including in super enhancers (*7, 8*). In physical space, super-resolution imaging techniques have recently described the spatial organization of the genome on the nanoscale. Together with chromosome conformation capture techniques these methods have shown that the genome is organized hierarchically by different processes on different length scales from nanometer (∼kilobase) scale clusters (*9*–*11*) to larger scale (∼100 kilobase scale) compartments (*12*–*15*). Due to limitations of statistical sample size and lack of genomic context in imaging experiments, mechanistic insights into how structures particularly on the scale of kilobases dynamically emerge, and how they interact with other biochemical processes involved in cell fate regulation, remain severely limited.

Here, we show how collective processes in physical space can be inferred from single-cell methylome sequencing measurements along the one-dimensional DNA sequence. We illustrate this approach using the example of the interplay between nanoscale topological changes in chromatin structure and the establishment of epigenetic marks (*de novo* methylation) during early mouse development. In early development, the formation of pluripotency is associated with the erasure of paternal and maternal DNA methylation (DNAme) in cytosines primarily in a CpG context (*16*). Upon exit from pluripotency which occurs around implantation of the embryo, upregulation of the *de novo* methyltransferase genes *Dnmt3a* and *b* leads to massive and rapid *de novo* methylation to a genome average of 80% per CpG (*17, 18*) which is associated with changes in chromatin structure (*9*). Here, combining single cell methylome data across this developmental period with a theoretical approach that transfers methods from theoretical physics (*21*) (field theory, renormalization group theory) to genomics we predict the spatial-temporal processes that underlie *de novo* DNA methylation. These processes are reflected in the spatial organization of chromatin and is confirmed by recent high-resolution microscopy studies and by orthogonal single cell multi-omics measurements. We also identify genomic elements that do not follow this general pattern, including gene body methylation patterns that precede silencing of pluripotency genes at the exit from pluripotency.

## Results

### De novo DNA methylation is a collective phenomenon involving spatial coordination across extended genomic domains

To develop our approach, we cultured mouse embryonic stem cells (mESCs) long-term in 2i culture conditions, where cells assume a naïve pluripotent state and DNAme is globally reduced. Cells were then released into serum conditions (Fig. 1B), where *Dnmt3a* and *b* genes are upregulated (*22*). The transition from 2i to serum conditions has been shown to recapitulate the epigenetic and transcriptional changes occurring during transition to formative and primed pluripotency in the embryo (*23*). After release into serum conditions we performed two complementary sets of experiments (Fig. 1C): 1) a whole-genome bisulfite-sequencing (BS-seq) time course of 31 time points over a period of 56 hours giving access to high-coverage information with high temporal resolution and 2) a single-cell NMT-sequencing (scNMT-seq) experiment of 288 mESCs with lower temporal resolution (0h, 24h and 48h) providing simultaneous information on the genomic distribution of DNAme and accessibility as well as on the transcriptome in single cells (*2*).

**Figure 1.**
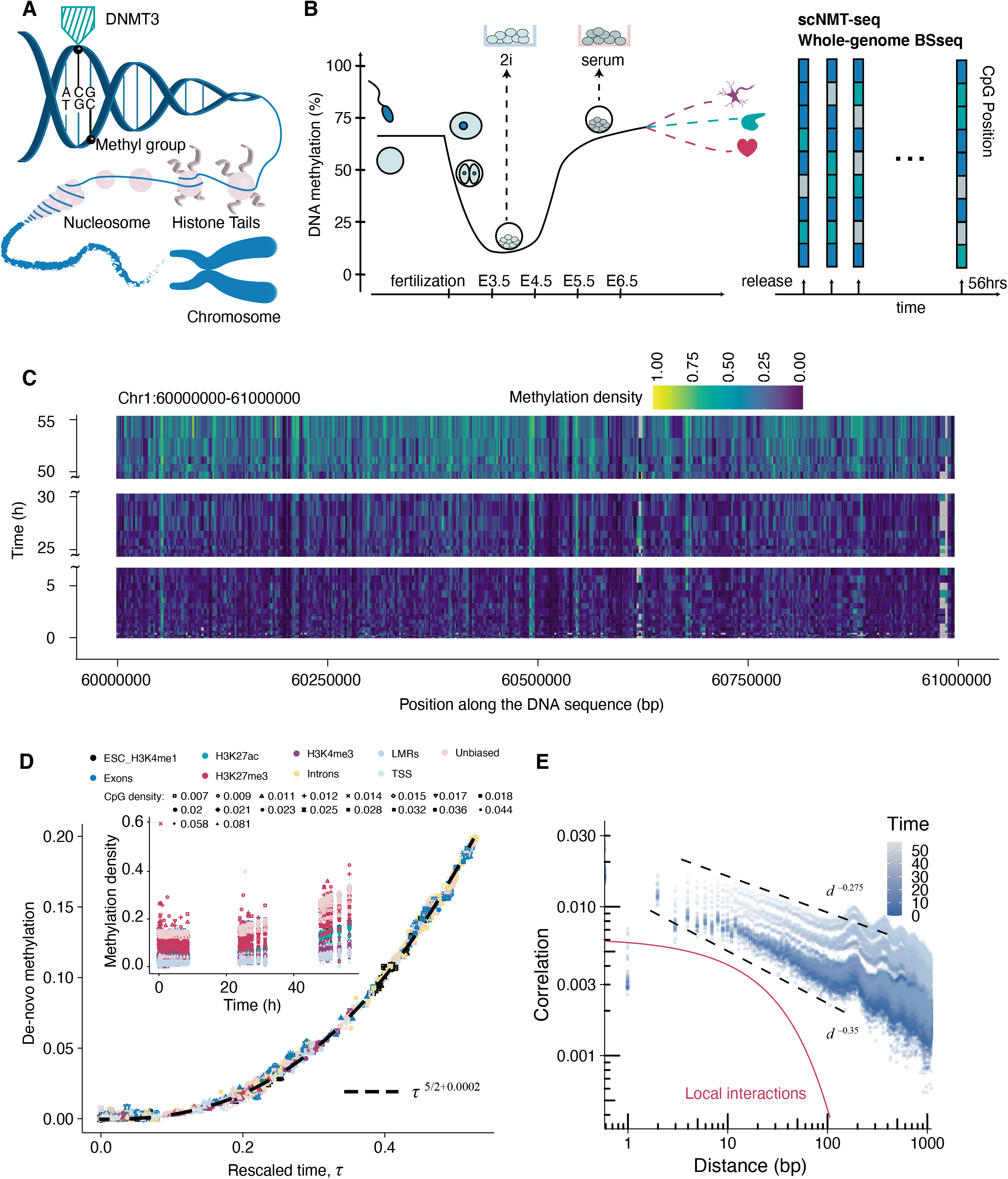
Scaling behaviour in *de novo* DNA methylation. (A) Schematic showing epigenetic processes involved in regulating cell fate. (B) DNA methylation dynamics in early development and 2i release experiment. (C) Heatmap showing DNA methylation levels across a genomic domain over time. (D) Inlay: Average DNA methylation levels increase at different rates in different regions with different functional annotation (colour) and CpG density (shape). Main plot: Rescaling time, the gain in average methylation follows a single power law with an exponent of 5/2 (dashed line). (E) Correlation functions show strong correlations in methylation marks. Dashed lines represent power laws with different exponents and the red line shows an exponential decay expected from local DNMT3 interactions.

To begin, we focused on the time evolution of DNAme levels during *de novo* methylation in the BS-seq experiment. The placement of methylation marks is thought to be regulated locally by a number of different factors, including CpG density, transcription and histone modifications (*24*). Indeed, we found that functionally distinct genomic regions (such as promoters, gene bodies etc.) of the genome acquired average DNA methylation levels at different rates (Fig. 1D, inlay). We were able, however, to rescale the time axis for each time series in such a way that all curves collapsed onto a single curve, a phenomenon referred to as scaling behaviour (Fig. 1E, Methods) (*25*). The emergence of scaling suggests that there is a generic mechanism of how DNAme is established genome-wide. Notably, the collapsed scaling curve follows almost perfectly a simple power law with a non-trivial exponent of 5/2 (Fig. 1E, Methods). The time evolution of average DNAme levels therefore is scale-invariant, i.e. its mathematical form does not change on time intervals of different lengths (self-similarity). The emergence of temporal scale-invariance and scaling behaviour are hallmarks of collective, self-organisation processes (*26*), suggesting that DNA methylation marks are established via a collective mechanism involving interacting DNMT3 enzyme molecules (*27, 28*). Importantly, if such interactions involve different genomic loci they might lead to the spatial coordination of *de novo* methylation (*26*).

To probe potential spatial coordination of *de novo* methylation we calculated the connected two-point correlation function as a summary statistic of the arrangement of methylation marks with respect to each other along the DNA sequence. Briefly, the value of the correlation function, also termed covariance, is proportional to the probability that two sites a given distance apart have the same methylation state and it captures correlations in the position of methylation marks independently of changes in the average, such as due to varying CpG density. As has been observed previously (*1*), we found that in addition to weak oscillations reflecting nucleosome positions, these correlations decay more slowly than expected by processes involving local interactions, following power laws over several orders of magnitude in sequence space (Fig. 1E). This reflects strong correlations of DNAme marks over distances of thousands of base pairs and which come about presumably by strong interactions between DNMT3 binding events over extended genomic domains. Taken together, these findings suggest that *de novo* methylation is a spatially coordinated collective phenomenon and hence provides a suitable and biologically interesting context to infer collective processes in physical space.

### Inference of de novo methylation kinetics in sequence space

To reveal the biological mechanisms underlying the interactions between DNMT3 binding events we developed a theoretical approach that allows inferring collective processes in the physical space of the nucleus from measurements along the one-dimensional sequence of the DNA (Fig. 2A). In contrast to typical hypothesis-driven approaches our framework deduces the kinetics from sequencing data. Briefly, we started by inferring the stochastic kinetics of *de novo* methylation in sequence space and then employed a dynamic geometric mapping between distances in sequence space and distances in physical space to derive the kinetics in physical space. This then allowed us to infer collective epigenetic phenomena in the physical space of the nucleus.

**Figure 2.**
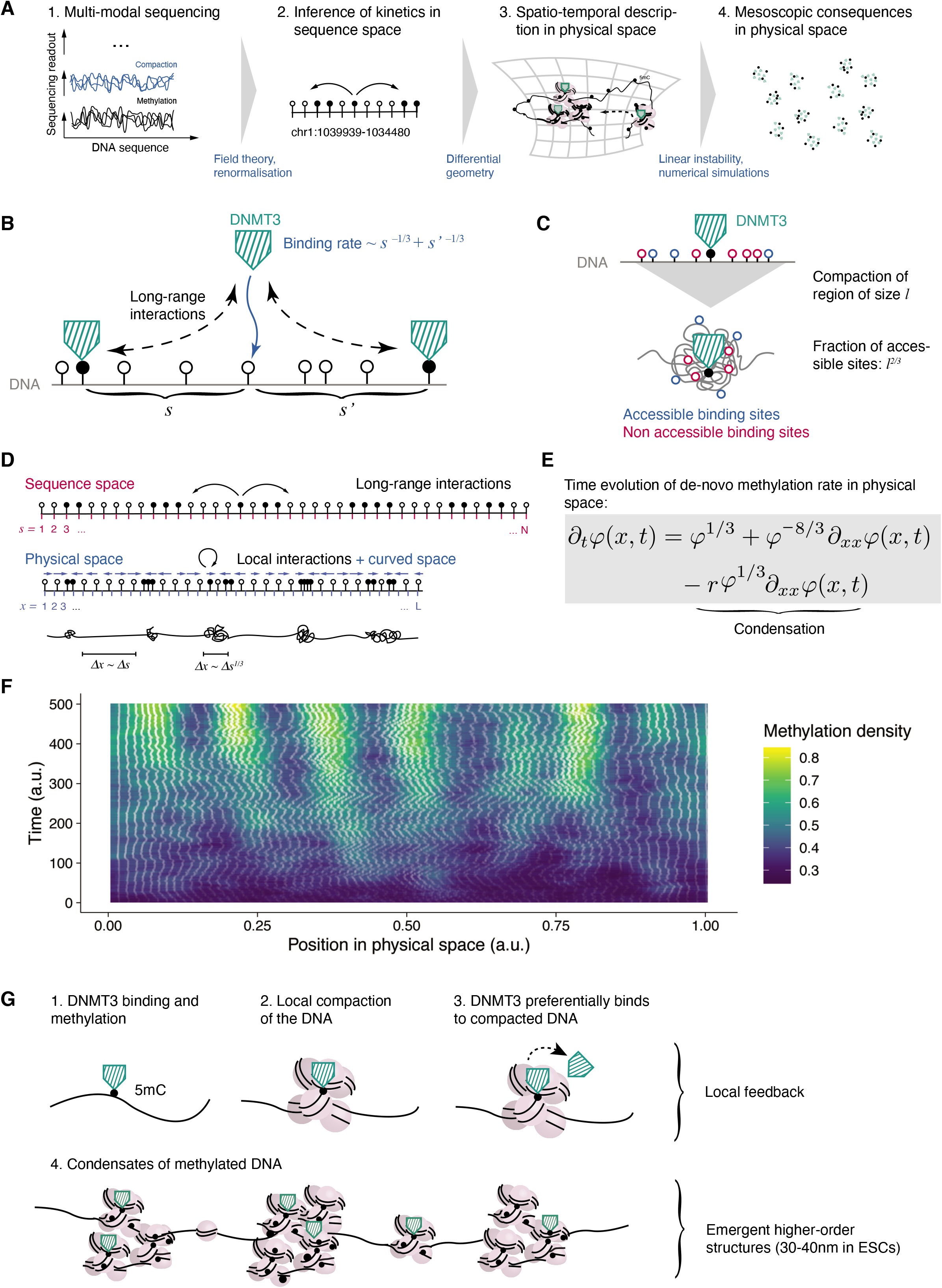
Inference of the spatio-temporal dynamics in physical space. (A) Summary of the theoretical approach. (B) Schematic illustrating the inferred quantification of interactions between methylation events in sequence space. (C) Geometric interpretation of the inferred interactions. (D) Illustration of the transformation from sequence space to physical space. (E) Equation describing the time evolution of the density of methylation marks, *φ*(*x, t*) in physical space. *r* is a parameter quantifying the effect of *de novo* methylation on DNA compaction (Supplemental Theory). (F) Stochastic simulation of the stochastic dynamics corresponding to the equation shown in (E), cf. Supplemental Theory. Colour denotes DNAme density and white lines denote trajectories of fixed positions in sequence space in physical space. (G) Summary of the inferred kinetic model and its mesoscopic consequences.

Specifically, we began by defining a mathematical framework (ansatz) for general stochastic enzyme kinetics incorporating 1) binding and unbinding of enzymes to the DNA and 2) chemical modifications of the DNA. Apart from these processes describing typical enzyme kinetics this framework also incorporates 3) general and unknown interactions of enzymes along the DNA sequence. These interactions between enzymes are described by an as yet unknown function quantifying how individual binding events influence each other (interaction kernel). Specifically, the interaction kernel quantifies the rate of binding to a locus as a function of the distance to other bound enzymes in its vicinity. It therefore captures processes specific to a given biological context and its inference is the central aim of the first step of our analysis. This ansatz therefore provides a general and flexible framework to describe enzyme-DNA interactions. Applied to DNA methylation, our mathematical framework entails the binding and unbinding of DNMT3 enzymes, and, if bound, the establishment of methylation marks. It can also contain active and passive demethylation processes, which in our case do not alter any of the results (Supplemental Theory).

Computer simulations of such stochastic systems with non-local interactions are computationally costly and converge poorly (*29*), such that they do not allow inferring the interaction kernel from sequencing data. By using methods originally developed in quantum theory and statistical physics (field theory, perturbation theory and renormalisation group theory) we were able to derive analytical solutions for a general class of interaction kernels, and how these interaction kernels relate to the statistics of the spatial and temporal evolution of DNAme marks. This in turn enabled us to infer the mathematical shape of interactions from static and dynamic sequencing data. Table 1 gives a summary of how different enzyme kinetics in sequence space are reflected in the statistics of linear sequencing measurements. Applied to DNA methylation, comparing the gain in average DNAme levels over time to the theoretical prediction then allowed us to infer the interaction kernel from the sequencing data. Specifically, we inferred that the binding rate of DNMT3 decreases with the distance to the nearest bound sites to the power of 1/3 (Fig. 2B). An alternative and equally feasible approach in our framework is to infer the interaction kernel from static, spatially resolved data, such as correlation functions. The inference of the interaction kernel fully determined the dynamics in sequence space.

**Table 1.**
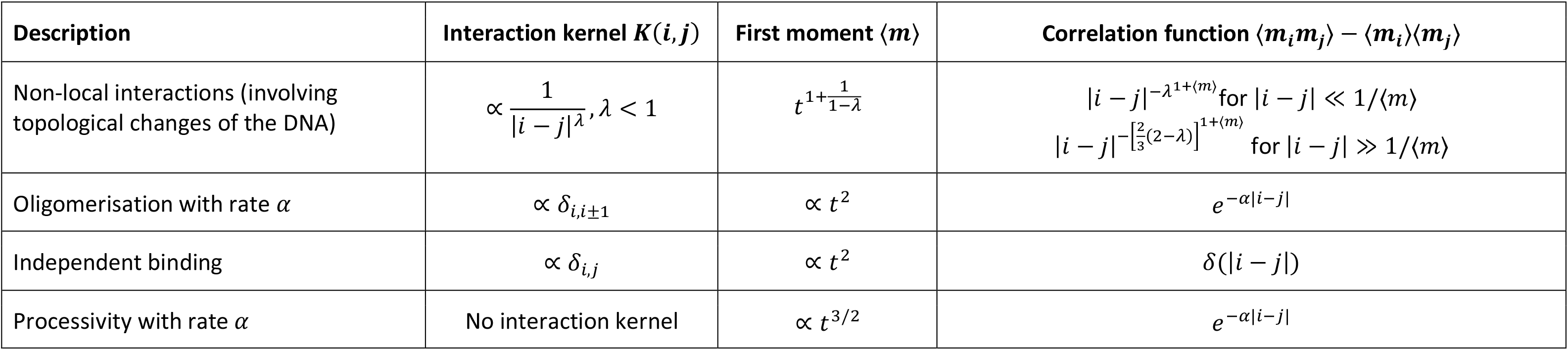
Statistical signatures of different classes of enzyme kinetics in sequencing measurements along the linear DNA sequence.

To interpret such interactions biologically, it is instructive to consider the total binding rate in the vicinity of size *l* of a bound site obtained by summing over all contributions in this region. Naively, the total binding rate in a genomic region should increase linearly with its length, *l*. In notable contrast, we found that only a fraction of sites is accessible for binding, *l*^2/3^. Indeed, *l*^2/3^ is the surface to volume ratio of an object with volume *l*, such that if a genomic region of *l* base pairs were compacted *l*^2/3^ base pairs would be accessible on the surface. Notably, the inferred interaction kernel describes the compaction of the DNA around methylated sites and the preferential binding of DNMT3 to compacted regions, resulting in positive feedback (Fig. 2C). This is fully consistent with biochemical studies which show that DNA methylation leads to tetra-nucleosome compaction *in vitro* (*30*) and with studies showing preferential binding of DNMT3 to compacted chromatin (*27, 31, 32*).

### Inference of higher-order chromatin dynamics in physical space

Having inferred the mathematical rules governing *de novo* DNAme along the one-dimensional sequence of the DNA, we then asked how such dynamics would manifest in the three-dimensional space of the cell nucleus. To this end, we calculated how small elements in sequence space change over time in physical space (Fig. 2D), leading to an effective mathematical description of the spatio-temporal dynamics in physical space (Fig. 2E). Such a mathematical description then allows predicting emergent (higher order) behaviour on larger scales in physical space: we found that the inferred dynamics resemble phase separation or spinodal decomposition processes (*26*), describing systems where two or more components – in our case methylated and nonmethylated DNA - separate spatially. Therefore, the local, constitutive feedback between DNAme and compaction (Fig. 2C) leads to the emergence of higher order chromatin structures, i.e. condensates of methylated DNA, in physical space importantly if the average DNAme level in a genomic region exceeds a threshold value (Fig. 3E, S1A). This is consistent with predictions from theoretical polymer models (*33, 34*). On larger genomic scales, we therefore predict the emergence of heterogeneously sized, local DNAme level dependent chromatin clusters. Their size is determined by a balance between DNAme-driven compaction and mechanical counteracting forces depending, for example, on the DNA rigidity. Although a prediction of the size of these clusters and the value of the threshold DNAme concentration relies on detailed quantitative knowledge of microscopic parameters, an order of magnitude estimate shows their size is in the order of 5000bp (40nm) (Supplemental Theory).

**Figure 3.**
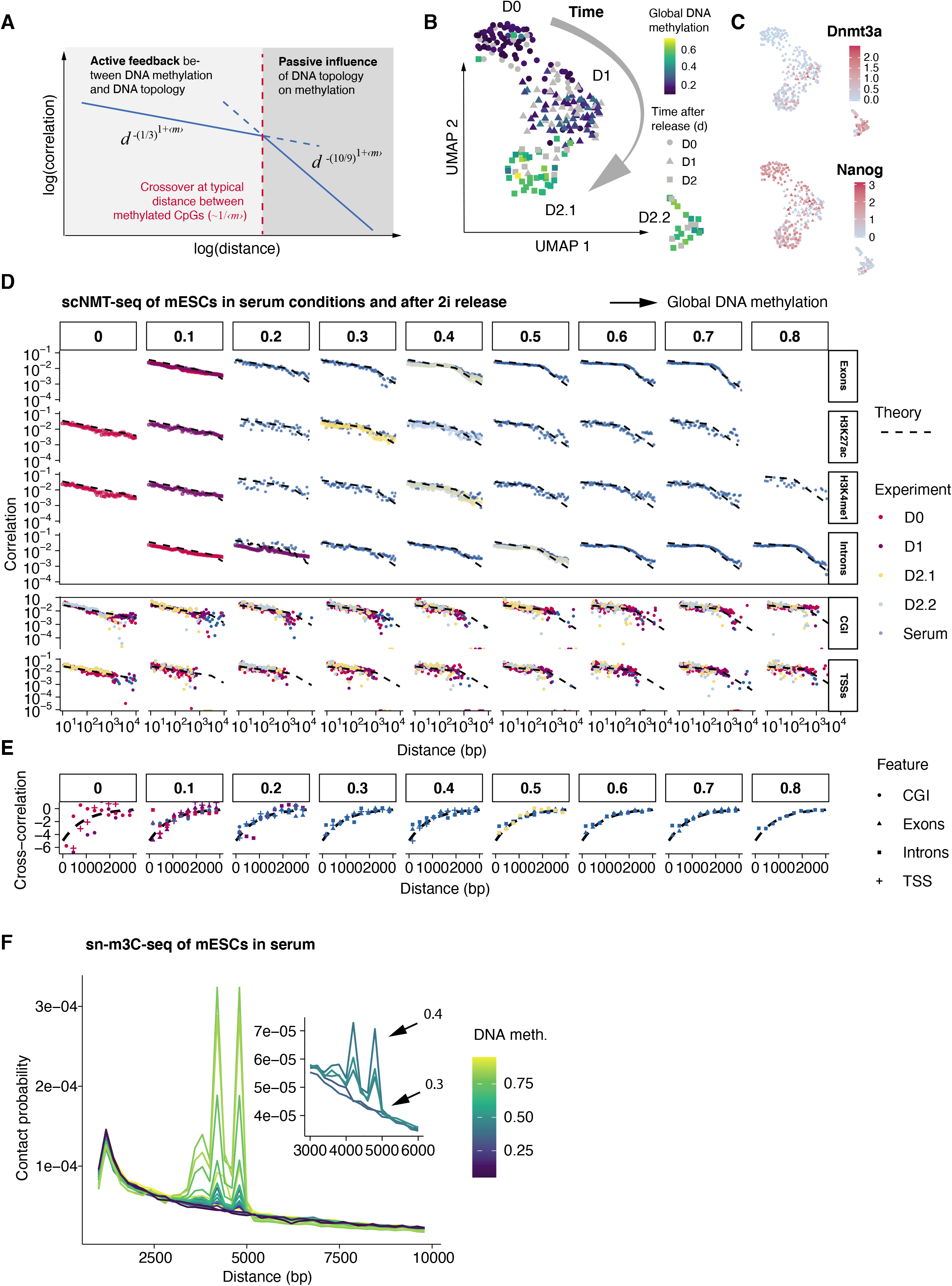
Test of model predictions. (A) Graphical summary of the processes determining the shape of correlations in DNA methylation. (B) Reduced-dimensional representation (UMAP) of gene expression profiles overlaid with average DNA methylation in each cell (colour). (C) UMAPs overlaid with expression levels of selected genes. (D) Prediction of correlation functions of DNA methylation in cells with different levels of global DNAme and empirical correlation functions from scNMT-seq experiments after 2i release and in serum conditions. (E) Prediction of the cross-correlation between DNAme and accessibility and empirical cross-correlations from scNMT-seq experiments after 2i release and in serum conditions. (F) Probability distribution of cis-contacts for windows of 100kbp size and varying average DNAme level in sn-m3C-seq data in serum ESCs. (G) Illustration showing the interpretation of the response function on different genomic scales. (H) Scale-dependent response function for scNMT-seq data from serum mESCs. Shaded regions represent standard errors. Values were corrected for biases stemming from varying bin sizes.

Remarkably, structures of strikingly similar size (roughly 12-40nm) have recently been described in super-resolution imaging experiments (*9*–*11*) and correlated with histone tail methylation and acetylation (*11*) and the presence of the H1 linker nucleosome (*9*). Notably, consistent with an association with DNAme these structures were found to be smaller in hypomethylated ESCs in 2i conditions compared to ESCs in serum conditions (*9*).

On the genome-scale, the inferred model predicts an increase of global DNAme according to a power law with an exponent of 5/2 as evidenced in the BS-seq time course (Supplementary Theory, Fig. 1D). Therefore, although the model describes processes on the nanometer or kilobase scale, these processes coordinate via long-range interactions (Fig. 2B) and determine the rate of increase of the global average DNAme level. Taken together, the feedback between chemical and topological modifications of the DNA leads to the emergence of higher order chromatin structures in highly methylated regions with genome-scale consequences on the time evolution of global DNAme levels. While our results do not rule out additional kinetics involved in DNA methylation, such as processivity (*35*), cooperativity (*36*), DNMT3 oligomerisation (*17*) or oscillatory dynamics (*22*), such processes are not statistically relevant on the genome-scale (Supplemental Theory). In addition, these processes give rise to quadratically increasing global DNAme levels and exponentially decaying correlation functions, but not the exponent of 5/2 and the long-range correlations in DNAme we observed (Table 1, Fig. 1D,E).

### Validating model prediction in sequence space using single-cell NMT-sequencing

To experimentally verify these findings, we sought to challenge them using experimental measurements not used for their inference. Having inferred the kinetics from purely temporal, averaged quantities (the time evolution of average methylation levels) we therefore asked whether the inferred kinetics could predict the spatial arrangement of methylation marks along the DNA sequence, as summarised in the correlation function, and resolved on the single-cell level by our scNMT-seq experiment. We found that *Dnmt3b* and *Dnmt3l* were expressed in most cells and *Dnm3a* was lowly expressed in a subset of cells 24h after release (Figs. S3C, S2B). As expected, global DNA methylation increased monotonically throughout the time course (Fig. S2C). In seeming contradiction to our prediction, on the global level accessibility increased slightly over time (Fig. S2C).

The inferred model gives insight into the processes governing the genomic distribution of DNAme marks on different length scales (Fig. 3A, Supplemental Theory): for distances, *d*, smaller than the typical distance between methylated sites the active interplay between *de novo* methylation and DNA compaction leads to a decay of the correlation function according to 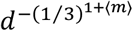, where ⟨*m*⟩ denotes the average methylation level. For longer-range distances these interactions become less relevant such that the distribution of DNAme is dominated by (passive) fluctuations in the conformation of the DNA. In this regime, the correlations between methylated CpGs decay faster, following 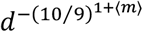. The position of the crossover between both regimes should therefore decrease with the average level of DNA methylation. Therefore, the genomic distribution of DNAme marks is dominated by different sources of biological fluctuations depending on the genomic length scale. This is reflected in different rates of decay of the correlation function on short and distant genomic length scales (Fig. 3A). We verified these analytical findings using stochastic simulations on a virtual DNA which resembled the statistics of CpG densities taken from the mouse genome (Fig. S2A). Comparing to our scNMT-seq experiment, we found that the distribution of DNAme sites along the DNA sequence indeed follows the predicted two-regime form for all studied genomic features (Fig. 3D). In addition, although the model does not have any free parameters, it quantitatively predicted the shape of correlation functions in a variety of genomic annotations excellently (Fig. 3D). The model also excellently predicted the spatial arrangement of DNA methylation in steady state serum cells across a range of global methylation levels (Fig. 3D).

To further challenge the mechanistic basis of the model we asked whether it was capable of predicting the local association between DNAme and DNA accessibility, which is summarised in the connected cross-correlation (or cross-covariance) between both quantities. Roughly, the cross-correlation function quantifies the correlation between DNAme and DNA accessibility at sites a given distance apart. We found that in agreement with the inferred model *de novo* methylation is associated with increased compaction locally in a region of roughly 1000bp as evidenced by the increasing strength of the cross-correlation between both quantities with time and DNAme level (Fig. S3A,B, for an analysis of the prediction of higher order structures see below). By including a negative feedback of DNAme on accessibility explicitly in our model in sequence space we predicted the interplay between DNAme and accessibility also quantitatively (Fig. 3E, Supplemental Theory). In summary, the model inferred by our approach is capable of accurately predicting the statistics of the genomic distribution of DNAme and their association with chromatin accessibility across a broad range of genomic regions.

### Validating model predictions in physical space using data from single-nucleus methyl-3C sequencing

To test if, as deduced in Fig. 2E,F, the local feedback between DNAme and compaction also leads to the formation of higher-order chromatin structures (methyl-condensates) on larger spatial scales with increasing levels of DNA methylation we reasoned that such condensates should be identifiable as an excess of mid-range physical contacts between pairs of genomic loci in highly methylated regions as measured in chromatin conformation capture experiments. We analysed single-nucleus methyl-3C sequencing data of mouse ESCs which allows locally correlating DNAme to chromatin structure in single cells (*37*). We tiled the mouse genome into windows of 100kb and, for each window calculated average DNAme levels and the probability distribution of cis contact distances. We found an abrupt increase in mid-range contacts between 3000bp and 5000bp (translating to roughly 30-40nm in diameter) for regions exceeding an average DNAme level of 40% (Fig. 3F), in agreement with our prediction of a phase transition and the emergence of DNAme associated chromatin structures. The sizes of these structure are again consistent with our theoretical estimate and with those estimated from super-resolution imaging studies (*9*–*11*).

### Identification of specific methylation pattern in pluripotency genes prior to their silencing

Having inferred and tested the kinetic rules governing *de novo* methylation in mESCs *in vitro* we then asked whether we could predict the establishment of 5mC marks during exit from pluripotency and early gastrulation *in vivo*. The model derived above describes a single mechanism establishing DNAme genome-wide. Although we were able to predict the distribution of DNAme across a wide range of genomic regions in sequence space with this model (Fig. 3D) such a mechanism alone cannot encode biological information in DNAme patterns beyond the binding affinity of DNMT3 enzymes to the DNA. Therefore, we expect that when cells become primed for differentiation from E5.5 and carry lineage-dependent DNAme patterns (*38*) additional mechanisms targeting DNA methylation must be in place. We reasoned that, by quantifying statistical patterns of deviations from our model describing generic, genome-wide DNAme dynamics (“null model”), we could identify genomic regions being specifically regulated by additional processes.

To address this, we analysed scNMT-seq data from mouse exit from pluripotency and initial cell fate decisions during gastrulation (*38*). As expected, the model predicted the distribution of DNAme marks in pluripotent cells at E4.5 (Fig. 4A). During later stages of development (E5.5-E7.5), when cells undergo cell fate transition changes, we observed systematic deviations between theory and experiment: while correlation functions still roughly followed the pattern of two distinct short and longer-distance spatial regimes (Fig. 3A) we found an enrichment in correlations in DNA methylation on a scale between 100 and 1000bp (Fig. S4A). To examine whether this pattern occurs genome-wide or is restricted to specific genomic regions we systematically quantified the difference between theory and experiment (residuals), normalised by the experimental standard error, for any distance between CpGs and different genomic annotations We found that the enrichment in correlated DNAme marks was specific to gene bodies (Fig. 4B, S4B), and in particular to genes silenced between E5.5 and E7.5, but not active genes (Fig. 4C). Absolute levels of DNAme in active and silenced genes differed only slightly (Fig. S4C) and therefore cannot fully explain these patterns.

**Figure 4.**
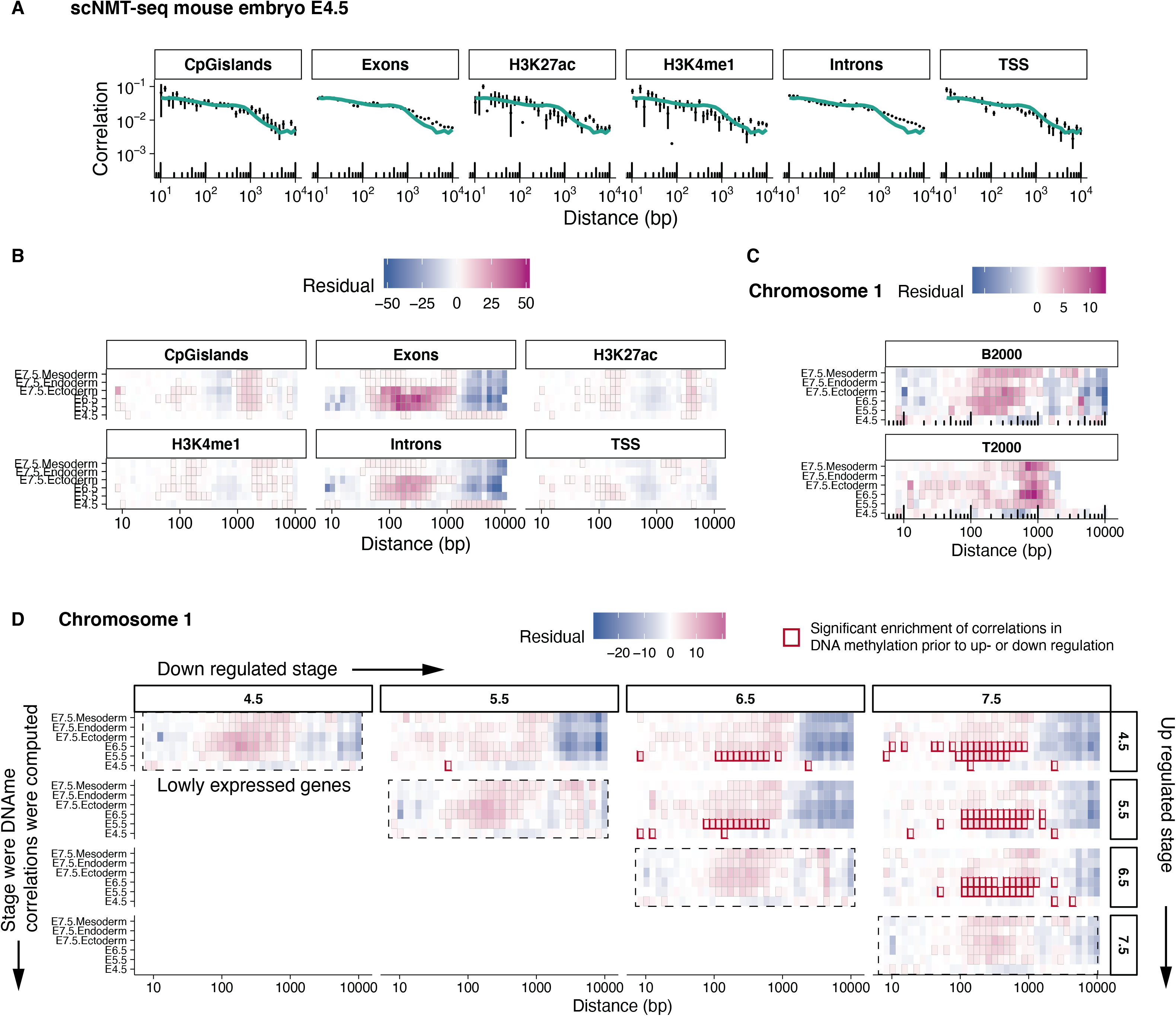
Identification of specifically regulated genomic regions in mouse development. (A) Theoretical and empirical correlation functions from scNMT-seq data of mouse embryos at E4.5. (B-D) Heatmaps showing differences between predicted and observed correlations in DNAme rescaled by the experimental standard error for (B) a set of functional annotations, (C) the gene bodies of the top and bottom 2000 expressed genes, and (D) for groups of genes that are differentially downregulated between pairs of embryonic stages. Pink regions signify an excess of correlation of the methylation state of pairs of CpGs at a given distance. Significant deviations are marked by black squares (p<0.05, t-test) and significant deviations preceding changes in gene expression are marked by red squares.

We then asked whether such a pattern is a consequence of gene silencing between E5.5 and E7.5, or whether it temporally precedes the silencing of genes during differentiation. To this end, we determined differentially expressed genes between each pair of embryonic stages and calculated for each set of genes the enrichment or depletion in spatial correlations between DNAme marks in all stages and lineages. We found that for silenced genes which are downregulated between a pair of embryonic stages these changes in DNAme patterns emerge up to two days before changes in the transcriptome appear, suggesting that these marks could play an instructive role by priming the genes for silencing during differentiation (Fig. 4D). By contrast, we identified the DNAme pattern characteristic for active genes (Fig. 4C, bottom) only after genes had been activated, but not before (Fig. S5A). We found that these patterns in particular apply to pluripotency genes (Fig. S5B, Table S1), but also to a set of silenced genes not annotated as pluripotency genes (Fig. S5C). While polycomb (H3K27me3) (*39*–*41*) or H3K9me3 (*42, 43*) pathways might be candidates for premarking silenced genes, further mechanistic studies will be necessary to elucidate the detailed molecular pathways behind this process. Taken together, our framework to infer collective epigenetic processes involved in *de novo* methylation allows identifying epigenetic patterns preceding transcriptional silencing during differentiation.

## Discussion

Using tools from theoretical physics in combination with multi-modal sequencing we demonstrated how measurements along the linear DNA sequence can inform about emergent processes in the physical space of the nucleus. Employing this framework in the context of *de novo* DNA methylation we found that feedback between chemical and topological processes leads to the formation of methyl-condensates on the scale of roughly 30-40nm in physical space. Although this process occurs on relatively small length scales it leads to coordination of de-novo methylation on larger genomic scales and ultimately has genome-wide consequences as evidenced by the emergence of scale-invariance in the time evolution of global DNA-methylation levels (Fig. 1D).

The condensates of methylated DNA reflect in size structures observed previously in super-resolution imaging experiments (*9*–*11*). The feedback between DNAme and compaction could be explained by a biophysical mechanism such as a charge-density driven phase transition in polyelectrolytes (*44*), or more complex pathways involving known chromatin modifiers such as MECP2 or HP1 and possibly nucleosome remodeling (*32, 45*). Super-resolution imaging studies have shown that the H1 nucleosome is necessary for the formation of similarly sized structures (*9*). Higher order structures of several hundred nanometers in size by contrast have been shown to be independent of DNAme (*12*–*15*). This is consistent with the lack of a global association between DNAme and compaction after 2i release (Fig. S2C) and highlights that different processes organize chromatin structure on different spatial scales. In the future, super-resolution imaging techniques as in (*9*–*11*) could be applied to DNMT3 knock out cells to independently validate our findings.

We derived these findings using a novel framework that allows for the inference of intermediary (mesoscopic) scale chromatin dynamics from linear sequencing measurements. This framework is based on a statistical and geometrical field theory which does not make assumptions about the details of molecular processes driving, for example, DNA compaction, such as nucleosomes. It is therefore sufficiently general to be easily applied and extended to parallel biological contexts involving enzyme-genome interactions. Our work emphasizes that analytical approaches originally developed in the context of quantum physics and for the first time applied to single-cell genomics in this work can solve problems in single-cell genomics that are intractable using more conventional computational approaches.

## Acknowledgements

We thank all members of the Reik and Rulands laboratories, S. Choubey, B. D. Simons, F. Piazza, and F. Jülicher for helpful discussions. A.P. is supported by a Sir Henry Wellcome Fellowship (215912/Z/19/Z). This project has received funding from the European Research Council (ERC) under the European Union’s Horizon 2020 research and innovation programme (grant agreement No. 950349). Research in the Reik laboratory is supported by the Biotechnology and Biological Sciences Research Council (BB/K010867/1) and the Wellcome Trust (095645/Z/11/Z).

## Author contributions

F.O., W.R. and S.R. conceived the project. T.L., S.J.C. and L.B. performed experiments. F.O. developed theory and performed modelling. F.O. and S.R. performed statistical analysis. F.O. and F.K. processed and managed sequencing data. F.O., S.R. and W.R. interpreted results and drafted the manuscript. All authors edited and approved the final manuscript.

## Competing interests

W.R. is a consultant and shareholder of Cambridge Epigenetix. The remaining authors declare no competing financial interests.

## Experimental Methods

### 2i release whole-genome bisulfite sequencing

For the 2i release experiment, cells were lysed by removing media from culture dishes and adding 200ul of RLT plus buffer (Qiagen) supplemented with 0.5mM 2-mercaptoethanol. In the first experiment, triplicate samples were collected at 31 time points from 0 h to 56 h 30 min. Further processing was performed identically to (*22*). Bisulfite sequencing (BS-seq) libraries were prepared from the total nucleic acid using the bulk-cell PBAT method previously described (Smallwood et al., 2014). Briefly, bisulfite conversion and purification was carried out using the EZ Methylation Direct MagPrep kit (Zymo), following the manufacturers’ instructions but with half volumes. Bisulfite converted DNA was eluted from MagBeads directly into 39ul of first strand synthesis reaction mastermix (1x Blue Buffer (Enzymatics), 0.4mM dNTP mix (Roche), 0.4uM 6NF preamp oligo (IDT) then heated to 65°C for 3 minutes and cooled on ice. 50U of klenow exo-(Enzymatics) was added and the mixture incubated on a thermocycler at 37°C for 30 minutes after slowly ramping from 4°C. Reactions were diluted to 100μl and 20U of exonuclease I (NEB) added and incubated at 37°C before purification using a 0.8:1 ratio of AMPure XP beads. Purified products were resuspended in 50μl of second strand mastermix (1x Blue Buffer (Enzymatics), 0.4mM dNTP mix (Roche), 0.4uM 6NR adaptor 2 oligo (IDT) then heated to 98°C for 2 minutes and cooled on ice. 50U of klenow exo-(Enzymatics) was added and the mixture incubated on a thermocycler at 37°C for 90 minutes after slowly ramping from 4°C. Second strand products were purified using a 0.8:1 ratio of AMPure XP beads and resuspended in 50μl of PCR mastermix (1x KAPA HiFi Readymix, 0.2uM PE1.0 primer, 0.2uM iTAG index primer) and amplified with 9 cycles. The final libraries were purified using a 0.8:1 volumetric ratio of AMPure XP beads before pooling and sequencing. All libraries were prepared in parallel with the pre-PCR purification steps carried out using a Bravo Workstation pipetting robot (Agilent Technologies). 9-12 libraries were sequenced as a multiplex on one Illumina HiSeq 2000 lane using 125bp paired-end read length.

### 2i release single-cell NMT sequencing

E14 mESC were cultured in 2i+LIF media consisting out of N2B27 medium, composed out of DMEM/F12 (Life Technologies, 12634010) and Neurobasal (Life Technologies, 21103049) in a volume/volume (v/v) ratio 1:1, 0.1 mM 2-mercaptoethanol, 2 mM L-Glutamine, 1:200 v/v N2 (Life Technologies, 17502048) and 1:100 v/v B27 supplement (Life Technologies, 17504044), supplemented with 3 µM CHIR99021, 1 µM PD0325901 and 20 ng/ml of leukemia inhibitory factor (LIF) (all Department of Biochemistry, University of Cambridge). mESCs were cultured on tissue culture plastic pre-coated with 0.1 % gelatine in H_2_O. Cultures were passaged when approaching confluence (every 2 days). All cells were cultured in a humidified incubator at 37 °C in 5 % CO_2_ and 20 % O_2_. mESCs were subject to routine mycoplasma testing using the MycoAlert testing kit (Lonza) and tested negative.

For induction for 2i-relase experiment, the cell been rinsed with PBS before the culture media was changed to serum + LIF medium consisting of DMEM (Life Technologies, 10566-016), containing 15% Fetal Bovine Serum (FBS) (Gibco, 10270106), 1x non-essential amino acids (NEAA) (Life Technologies, 11140050), 0.1 mM 2-mercaptoethanol (Life Technologies, 31350-010), 2 mM L-Glutamine (Life Technologies, 25030-024), supplemented with 20 ng/ml of LIF (Department of Biochemistry, University of Cambridge).

0h, 24h, and 48h after induction of 2i-release the cells been dissociated into single cells using accutase before flow sorting (BD Influx) into 96-well plate containing 2.5μl of methylase reaction buffer (1 × M.CviPI Reaction buffer (NEB), 2 U M.CviPI (NEB), 160 μM S-adenosylmethionine (NEB), 1 U μl^−1^ RNasein (Promega), 0.1% IGEPAL CA-630 (Sigma)). Samples were incubated for 15 min at 37 °C to methylate accessible chromatin before the reaction was stopped with the addition of RLT plus buffer (Qiagen) and samples frozen down and stored at −80 °C before processing. All downstream library preparation steps been performed as previously described by Clark et al. (Clark et al. Nature Communications, 2018).

All sequencing was carried out on HiSeq instrument. BS-seq libraries were sequenced in 96-plex pools using 125-bp paired-end reads. RNA-seq libraries were pooled as 96-plex or 192-plex pools and sequenced using 75-bp paired-end reads.

### Quantification and statistical analysis

#### Processing of whole-genome bisulfite sequencing data

Whole genome bisulfite sequencing data was processed identically to(*22*). Raw sequence reads were trimmed to remove both poor-quality calls and adapters using Trim Galore (v0.4.1, www.bioinformatics.babraham.ac.uk/projects/trim_galore/, Cutadapt version 1.8.1, parameters: -- paired) (Martin, 2011). Trimmed reads were first aligned to the mouse genome in paired-end mode to be able to use overlapping parts of the reads only once while writing out unmapped singleton reads; in a second step remaining singleton reads were aligned in single-end mode. Alignments were carried out with Bismark v0.14.4 (Krueger and Andrews, 2011) with the following set of parameters: a) paired-end mode: --pbat; b) single-end mode for Read 1: --pbat; c) single-end mode for Read 2: defaults. Reads were then deduplicated with deduplicate_bismark selecting a random alignment for position that were covered more than once. CpG methylation calls were extracted from the deduplicated mapping output ignoring the first 6 bp of each read (corresponding to the 6N random priming oligos) using the Bismark methylation extractor (v0.14.4) with the following parameters: a) paired-end mode: --ignore 6 -- ignore_r2 6; b) single-end mode: --ignore 6. SeqMonk version 0.32 was used to compute methylation rates and coverage in annotation genomic regions. To QC BS-seq data, pairwise Pearson correlation coefficients were calculated using methylation levels averaged over 10kb tiles. Replicates within the same time point were on average more highly correlated than between time points (r=0.885 versus 0.866). For subsequent analyses, replicates were merged.

Further statistical analysis was performed by custom scripts in R. For Figure 1D, we calculated average methylation levels for a given set of genomic regions defined by their functional annotation and average CpG density using the “Bisulfite methylation over feature” pipeline in Seqmonk. To be able to identify the functional form of average methylation over time only feature sets that have genome-wide more than 1500 reads at a given time point are shown. Averages over genomic regions were weighted by the average number of reads per CpG.

To collapse the time series onto a scaling form, we made a scaling ansatz of the form ⟨*m*⟩ = *a* + *b t*^5/2^ and determined *a* and *b* using a nonlinear least squares estimate as implemented in the R function nls. With this, the rescaled time, *τ*, was defined as *τ* = *t* ∗ *b*^2/5^. The exponent was estimated using nonlinear least squares. To verify the robustness of the exponent with respect to log transformation of both axes we estimated the exponent for different values of an offset parameter, *c*, such that the rescaled average DNA methylation reads ⟨*μ*⟩ = *c* + *τ*^*γ*^ and all values of the time course are positive. We found that under these transformations the estimation of the exponent was robust.

### Processing of scNMT-seq 2i release data

#### BS-seq

Cells with more than 10^5.5^ reads, less than 15% CHH methylation and a mapping efficiency larger than 10% were kept for downstream analysis. Following (*46*) average DNA methylation was calculated as

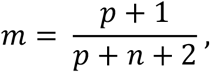

where *p* and *n* signify the number of positive or negative reads in a given genomic interval, respectively.

#### RNA-seq

226 cells with mitochondrial RNA < 0.15%, > 200000 reads and > 2000 detected genes were kept for downstream analysis. Reads were log normalised using the LogNormalise function of the Seurat package version 3.2.0 with standard parameters. For dimensionality reduction, the top 1000 most highly variable genes were selected and a principal component analysis with default parameters of the Seurat package was performed. Uniform Manifold Approximation was performed on the 15 principle components with the highest variance and with a minimum distance of 0.2.

### Processing of sn-m3C-seq data

Following (*37*) we retained cells with more than 5000 cis contacts at distances longer than 10000bp and more than 100000 covered CpGs. We tiled the genome into windows of 100kbp and, for each tile, calculated average DNAme and cis contact histograms with respect to the genomic distance. We then pooled these histograms for genomic windows of similar DNAme levels and normalized by the total number of cis contacts.

### Processing of scNMT-seq data of mESCs (serum)

Data was processed identically to (*2*).

### Processing of scNMT-seq embryo data

#### BS-seq

Data was processed identically to (*38*). Genome-wide correlation and cross-correlation functions were computed by dividing samples with respect to the stage (E4.5, E5.5, E6.5) and lineage (E7.5 Mesoderm, Endoderm, Ectoderm).

#### RNA-Seq

Cells which had a percentage of mitochondrial RNA <0.15%, nCount_RNA>1e5 and more than 2500 genes with at least one read were kept for downstream analysis. Normalisation was performed using the function LogNormalize from the Seurat package (version 3.2.9). The the least and most highly expressed genes were determined based on their log-normalised expression value. Differentially expressed genes between pairs of stages was determined using a t-test. To keep the sample size comparable the 5000 most significant genes based on p-value were selected for further analysis. Correlations functions for a given set of genes were computed by first obtaining the coordinates of the corresponding gene bodies using biomart 2.44.1, then computing correlation functions for each gene and finall averaging over all the genes in a given stage or lineage.

### Correlation and cross-correlation functions

Connected correlation functions for a given distance, *j*, were defined as

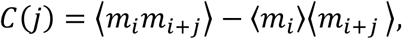

where *m*_*i*_ and *m*_*I*+*j*_ are the methylation states of a CpG at position *i* and *i* + *j*respectively. The average ⟨… ⟩ is performed over all pairs of CpGs that are a distance *j* apart and over all the samples as described below. To compute (cross-) correlation functions we sought to group statistically similar samples. To this end, we grouped cells with similar global levels of DNA methylation and average correlations in a given annotation over these cells. For promoters and CpG islands, which show bimodal average DNA methylation levels or where methylation levels are less strictly correlated to global DNA methylation in a cell(*22*), in a given cell we averaged over all regions with similar DNA methylation level and then averaged over all cells. For the sc-NMTseq embryo data samples were grouped by embryonic stage and lineage since within these groups variance in global DNA methylation levels is low compared to serum conditions. Analogously, connected cross-correlation functions were defined by,

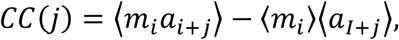

where *a*_*i*+*j*_ is the accessibility (GpC methylation) state at position *i* + *j*. The average ⟨… ⟩ was performed in the same way as for correlation functions. All correlation and cross correlation are normalized such that the area underlying the curve, which is in the case of correlation functions is dependent on sequencing coverage, is equal to 1. To predict the cross-correlation function we estimated the length scale of local chromatin compaction using R’s nls function.

### Conversion between genomic and physical distances

To convert sizes of compact chromatin structures with radius *r* to genomic distances in base pairs, *n*, and vice versa we, in a rough approximation, equated the spherical volume in physical space, 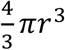 and the volume occupied by *N* nucleosomes,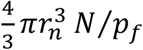. Here, *r*_*n*_ ≈ 5.5*mm* is the radius of a single nucleosome and *p*_*f*_ = 0.64 is the random packing fraction. Genomic distances were obtained by solving for *N* and using that a single nucleosome corresponds to roughly 200bp in sequence space.

