## Supplementary Materials for "Inference of emergent spatio-temporal processes from single-cell sequencing reveals feedback between *de novo* DNA methylation and chromatin condensates"

**This PDF file contains:**

Supplementary Figure legends

Supplementary Figures S1-S5

Supplementary Tables S1-S2

Supplementary Theory

**Supplementary figure legends**

**Figure S1. (A)** Linear stability analysis considers the rate of growth of small perturbations with a given wave number, $k$ (spatial frequency). It shows that for sufficiently high levels of DNAme the equation presented in Fig. 2E leads to the growth of perturbations with a finite wave number, and therefore the formation of mesoscopic patterns in the methylation density.

**Figure S2. (A)** Comparison of our analytical prediction with a stochastic simulation performed on a virtual DNA statistically resembling the distribution of CpGs in the mouse genome (Mean +/- SEM, Supplemental Theory). **(B)** Average accessibility (top) and DNA methylation (bottom) over the time course of the 2i release scNMT-seq experiment. Each dot represents a cell, lower and upper hinges corresponds to the first and third quartile and whiskers extend to 1.5 times the inter quartile range. **(C)** Log-normalised expression of a set of genes shown on a UMAP projection.

**Figure S3.** Making an ansatz $c(x) = a\, e^{-\frac{x}{b}}$ for the cross-correlation function we estimated **(A)** the strength, $b$, and **(B)** the length scale, $a$, of the cross-correlation between DNA methylation and accessibility using nonlinear least squares fitting. Error bars denote standard errors of the estimates.

**Figure S4. (A)** Empirical and simulated correlation functions of DNA methylation for different stages of embryonic development. Error bars represent standard errors and arrows indicate an enrichment of empirical correlations with respect to the model predictions. **(B)** Difference between simulated and empirical correlation function (residual) rescaled by the empirical standard error shown separately for each chromosome. Correlation functions were calculated for gene bodies corresponding to the bottom (left) and top (right) 2000 expressed genes. **(C)** Boxplot showing average gene body methylation for both groups of genes.

**Figure S5. (A)** Heatmaps showing differences between predicted and observed correlations in DNAme rescaled by the experimental standard error for groups of genes that are differentially up regulated between pairs of embryonic stages, as in Fig. 4D. Significant deviations are marked by black squares ($p<0.05$, t-test) and significant deviations preceding changes in gene expression are marked by red squares. The same heatmaps are shown for (B) gene bodies of pluripotency genes, and (C) gene bodies of the top and bottom 2000 expressed genes excluding pluripotency genes.

**Figure S1**

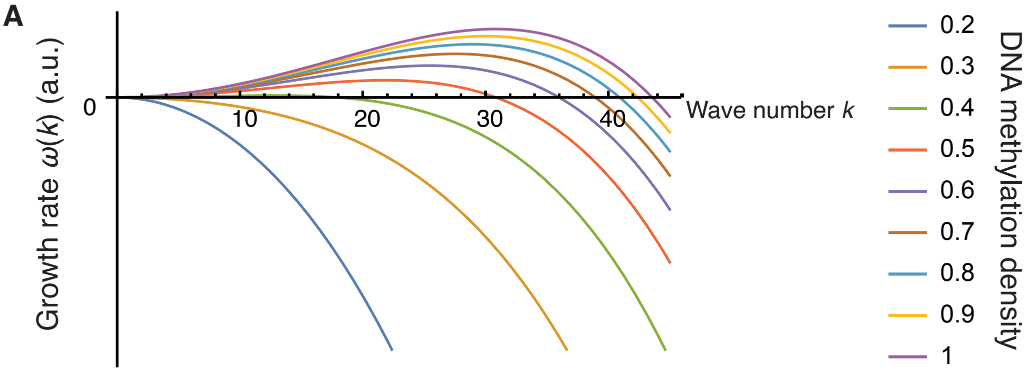
A
Growth rate ω(k) (a.u.)
0
10
20
30
40
Wave number k
0.2
0.3
0.4
0.5
0.6
0.7
0.8
0.9
1
DNA methylation density

**Figure S2**

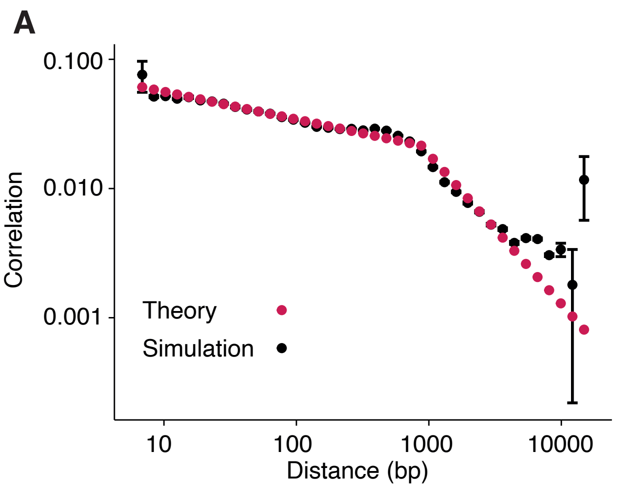
A
Correlation
0.100
0.010
0.001
Theory
Simulation
10
100
1000
10000
Distance (bp)

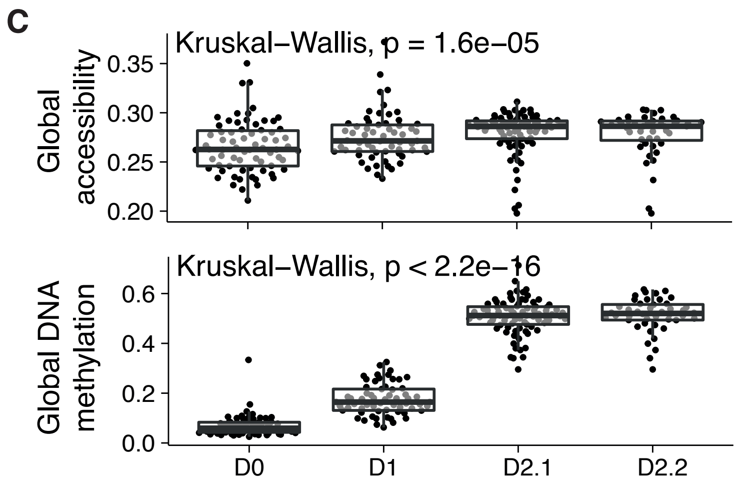
C
Global accessibility
Kruskal–Wallis, p = 1.6e–05
0.35
0.30
0.25
0.20
Global DNA methylation
Kruskal–Wallis, p < 2.2e–16
0.6
0.4
0.2
0.0
D0
D1
D2.1
D2.2

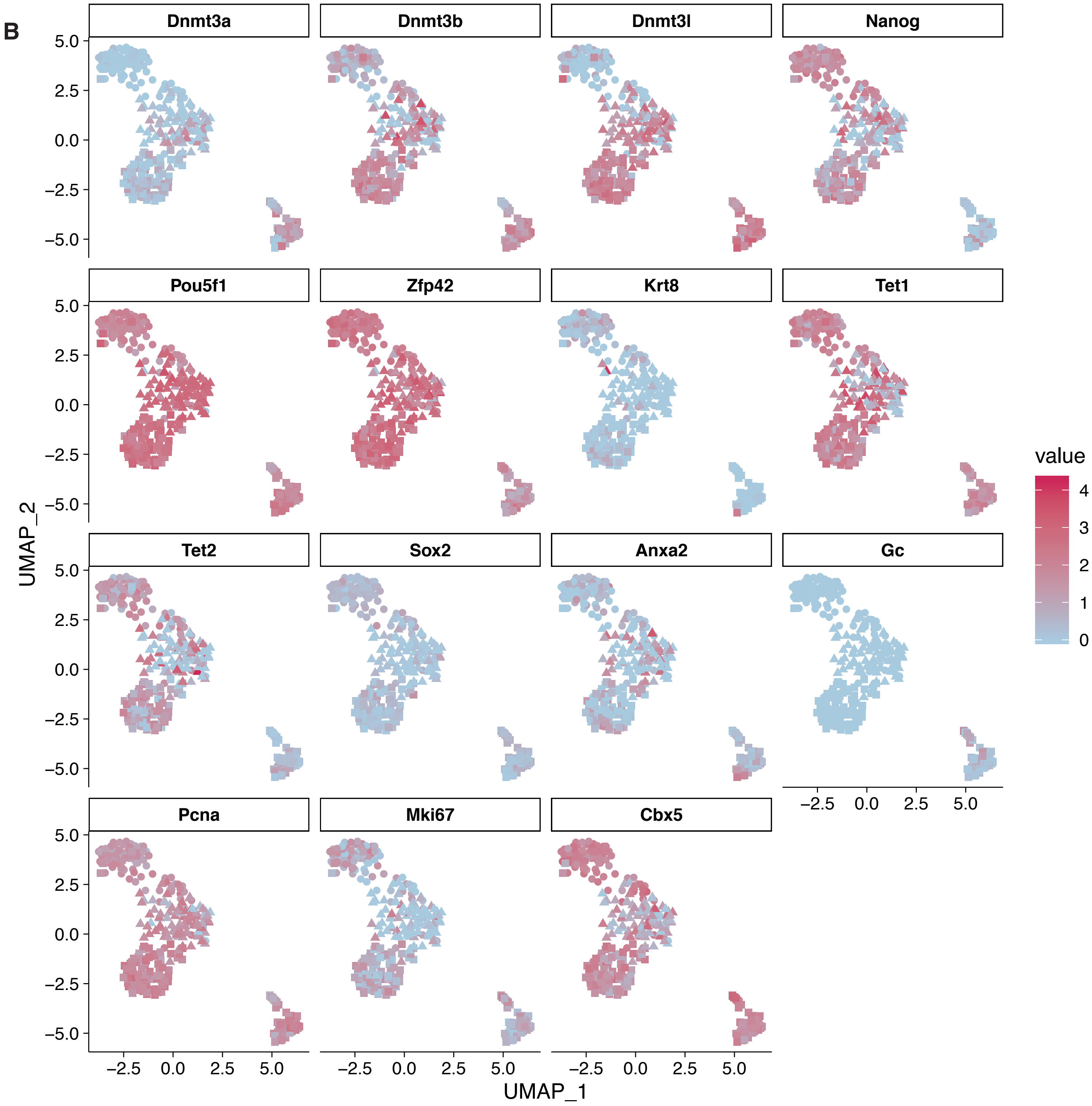
B
Dnmt3a
Dnmt3b
Dnmt3l
Nanog
Pou5f1
Zfp42
Krt8
Tet1
Tet2
Sox2
Anxa2
Gc
Pcna
Mki67
Cbx5
UMAP_2
UMAP_1
5.0
2.5
0.0
–2.5
–5.0
value
4
3
2
1
0

### Figure S3

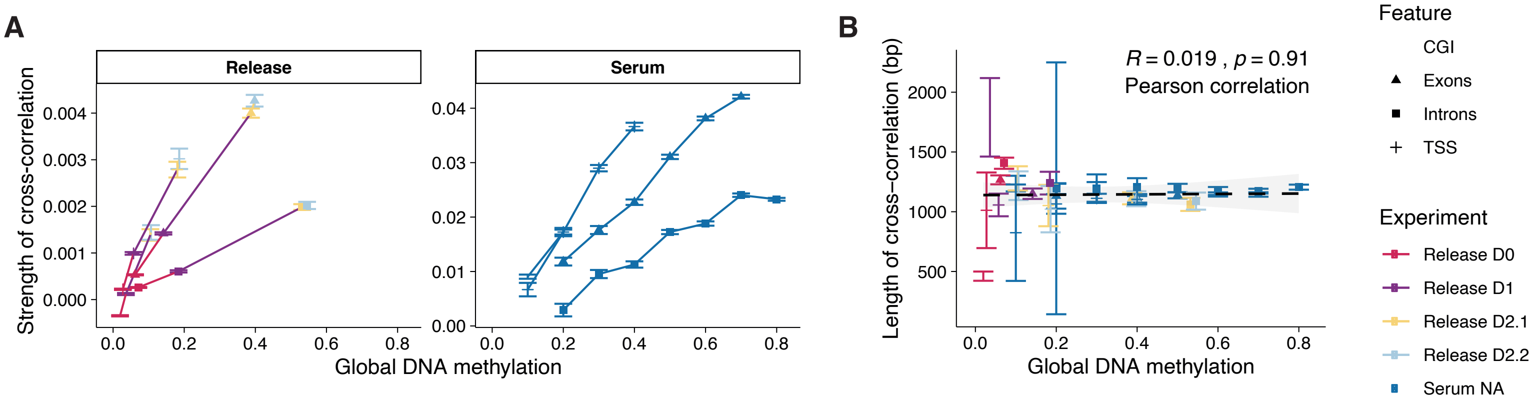
A
Release
Serum
Strength of cross-correlation
0.000
0.001
0.002
0.003
0.004
0.00
0.01
0.02
0.03
0.04
0.0
0.2
0.4
0.6
0.8
Global DNA methylation
B
R = 0.019 , p = 0.91
Pearson correlation
Length of cross-correlation (bp)
500
1000
1500
2000
Feature
CGI
Exons
Introns
TSS
Experiment
Release D0
Release D1
Release D2.1
Release D2.2
Serum NA

#### Figure S4

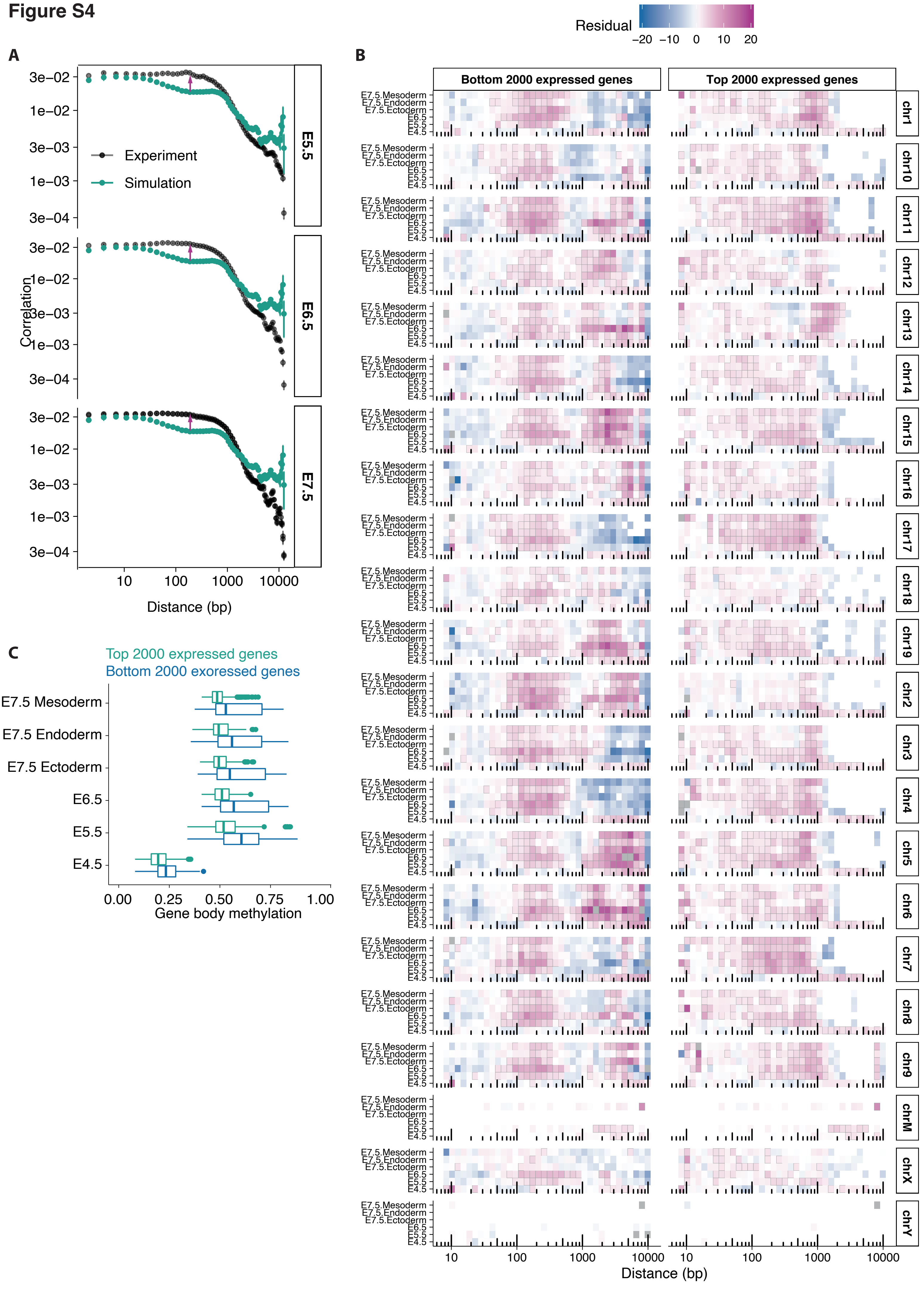
A
B
C
Residual
−20 −10 0 10 20
Bottom 2000 expressed genes
Top 2000 expressed genes
Experiment
Simulation
Correlation
E5.5
E6.5
E7.5
Distance (bp)
Top 2000 expressed genes
Bottom 2000 exoressed genes
E7.5 Mesoderm
E7.5 Endoderm
E7.5 Ectoderm
E6.5
E5.5
E4.5
Gene body methylation
chr1
chr10
chr11
chr12
chr13
chr14
chr15
chr16
chr17
chr18
chr19
chr2
chr3
chr4
chr5
chr6
chr7
chr8
chr9
chrM
chrX
chrY

### Figure S5

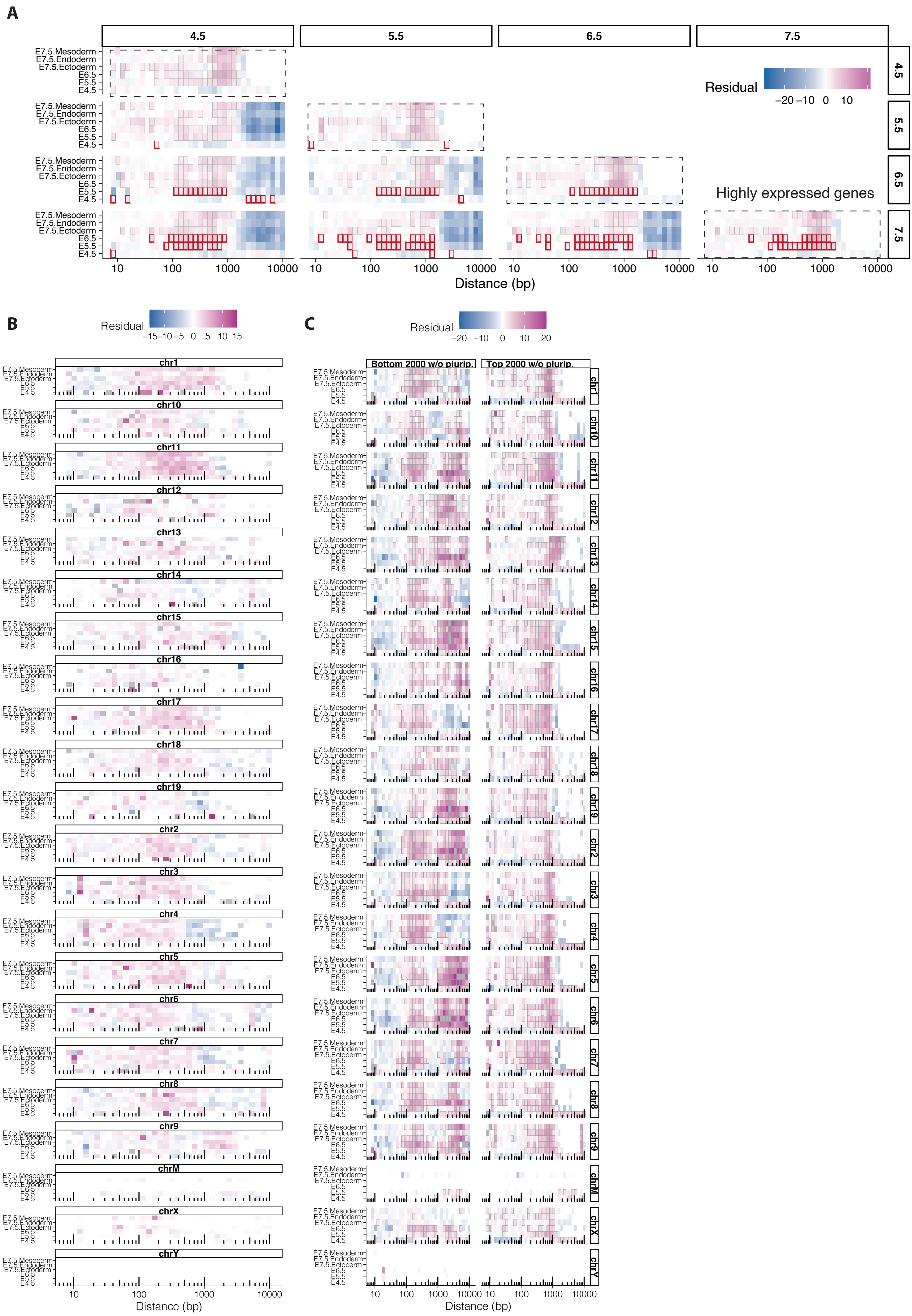
A
4.5
5.5
6.5
7.5
E7.5.Mesoderm
E7.5.Endoderm
E7.5.Ectoderm
E6.5
E5.5
E4.5
Residual
−20 −10 0 10
Highly expressed genes
10 100 1000 10000
Distance (bp)
B
Residual
−15−10 −5 0 5 10 15
chr1
chr10
chr11
chr12
chr13
chr14
chr15
chr16
chr17
chr18
chr19
chr2
chr3
chr4
chr5
chr6
chr7
chr8
chr9
chrM
chrX
chrY
10 100 1000 10000
Distance (bp)
C
Residual
−20 −10 0 10 20
Bottom 2000 w/o plurip.
Top 2000 w/o plurip.
10 100 1000 10000
Distance (bp)

**Table S1**

| Description | Reference |
| --- | --- |
| Primed ESC Dnmt3a/b ChIP-seq | Buecker et al (2014) Cell Stem Cell 14, 838-853 |
| Primed ESC H3K27ac ChIP-seq | Creyghton et al (2010) Proceedings of the National Academy of Sciences 107, 21931-21936 |
| Primed ESC H3K27me3 ChIP-seq | Creyghton et al (2010) Proceedings of the National Academy of Sciences 107, 21931-21936 |
| Primed ESC H3K4me3 ChIP-seq | Creyghton et al (2010) Proceedings of the National Academy of Sciences 107, 21931-21936 |
| Primed ESC p300 ChIP-seq | Creyghton et al (2010) Proceedings of the National Academy of Sciences 107, 21931-21936 |
| CpG Islands | Illingworth et al (2010) PLoS Genetics 6, e1001134 |
| Primed ESC low methylated regions (LMRs) | Stadler et al (2011) Nature 480, 490-495 |
| Mouse embryo scNMT-seq | Clark et al (2018) Nature Communications 9, 781 |
| Primed ESC sn-m3C-seq | Li et al (2019) Nat. Methods, 1-3 |
| List of pluripotency genes | Müller (2008) Nature 455(7211): 401–405 and Gerrard et al. (2016) eLife ;5:e15657 |
| Polycomb target genes | Bracken et al (2006) Genes & Dev. 20: 1123-1136 and Liu et al (2010) Genomics , 96,1, 17-26 |

**Table S2**

| p_val | avg_logFC | pct.1 | pct.2 | p_val_adj | cluster | gene |
|---|---|---|---|---|---|---|
| 9,16E-56 | 0,899753628 | 1 | 0,055 | 2,14E-51 | 3 | Gm42047 |
| 1,32E-50 | 0,804170057 | 0,957 | 0,044 | 3,10E-46 | 3 | Eif2s3y |
| 4,03E-50 | 1,122962524 | 1 | 0,088 | 9,44E-46 | 3 | Hprt |
| 6,40E-32 | 0,711607419 | 1 | 0,283 | 1,50E-27 | 3 | Pdpn |
| 1,12E-27 | 1,424763504 | 1 | 1 | 2,63E-23 | 3 | Rpl34 |
| 1,44E-27 | 1,313340228 | 1 | 0,904 | 3,38E-23 | 3 | Rpl29 |
| 3,65E-27 | 0,732180673 | 1 | 0,669 | 8,55E-23 | 3 | Ubc |
| 4,84E-27 | 0,904191457 | 1 | 0,996 | 1,13E-22 | 3 | Rpl17 |
| 1,07E-26 | 1,042858284 | 1 | 1 | 2,52E-22 | 3 | Rpl23 |
| 3,07E-26 | 1,07402312 | 1 | 0,989 | 7,18E-22 | 3 | Hmgn2 |
| 4,26E-26 | 0,838048523 | 1 | 1 | 9,97E-22 | 3 | Gm9794 |
| 4,50E-26 | 1,343934315 | 1 | 1 | 1,05E-21 | 3 | Ppia |
| 5,60E-26 | 1,327713615 | 1 | 1 | 1,31E-21 | 3 | Ftl1 |
| 1,46E-25 | 1,554419831 | 1 | 1 | 3,43E-21 | 3 | Gm15772 |
| 7,75E-25 | 1,308155898 | 1 | 1 | 1,82E-20 | 3 | Rps27 |
| 9,77E-25 | 1,012169149 | 1 | 1 | 2,29E-20 | 3 | Cbx1 |
| 1,59E-24 | 0,734645463 | 0,979 | 0,382 | 3,72E-20 | 3 | Cyct |
| 1,91E-24 | 0,970958446 | 1 | 1 | 4,48E-20 | 3 | Ncl |
| 2,45E-24 | 0,796591313 | 1 | 1 | 5,74E-20 | 3 | Gm14539 |
| 5,58E-24 | 1,169349184 | 1 | 1 | 1,31E-19 | 3 | Rpl38 |
| 5,88E-24 | 0,967594275 | 1 | 1 | 1,38E-19 | 3 | Npm1 |
| 1,10E-23 | 0,670186992 | 1 | 1 | 2,58E-19 | 3 | Gm12191 |
| 1,18E-23 | 0,786323236 | 1 | 1 | 2,76E-19 | 3 | Ywhae |
| 1,40E-23 | 0,793612912 | 1 | 0,978 | 3,28E-19 | 3 | Psmb5 |
| 2,44E-23 | 0,721999882 | 1 | 1 | 5,71E-19 | 3 | Gm9118 |
| 3,15E-23 | 0,941807469 | 1 | 1 | 7,39E-19 | 3 | Rps3a1 |
| 1,02E-22 | 0,906314403 | 1 | 1 | 2,40E-18 | 3 | Rplp1 |
| 1,51E-22 | 1,046037806 | 1 | 1 | 3,54E-18 | 3 | Rpl41 |
| 1,79E-22 | 0,853754027 | 1 | 1 | 4,19E-18 | 3 | Rps27a |
| 1,88E-22 | 0,854840754 | 1 | 1 | 4,41E-18 | 3 | Rps13 |
| 3,07E-22 | 0,700194215 | 1 | 1 | 7,19E-18 | 3 | Rpl35a |
| 3,14E-22 | 0,723939487 | 1 | 0,555 | 7,35E-18 | 3 | ENSMUSG00000116673 |
| 6,51E-22 | 0,807340211 | 1 | 1 | 1,52E-17 | 3 | Rps26 |
| 7,20E-22 | 0,844465998 | 1 | 0,989 | 1,68E-17 | 3 | Mrps24 |
| 1,20E-21 | 0,825421192 | 1 | 0,993 | 2,82E-17 | 3 | Tpt1 |
| 1,79E-21 | 0,907360952 | 1 | 0,996 | 4,19E-17 | 3 | Sod1 |
| 2,57E-21 | 0,830493055 | 1 | 0,794 | 6,02E-17 | 3 | Esd |
| 2,74E-21 | 0,86530211 | 1 | 0,996 | 6,42E-17 | 3 | Gm14681 |
| 7,30E-21 | 0,67105426 | 1 | 1 | 1,71E-16 | 3 | Rps7 |
| 7,79E-21 | 1,008312807 | 1 | 1 | 1,82E-16 | 3 | Uqcr10 |
| 8,70E-21 | 0,798061014 | 1 | 0,956 | 2,04E-16 | 3 | Polr2k |
| 1,64E-20 | 0,655660755 | 1 | 1 | 3,83E-16 | 3 | Pdcd5 |
| 2,15E-20 | 0,755304057 | 1 | 1 | 5,04E-16 | 3 | Rps12 |
| 2,35E-20 | 0,79230419 | 1 | 0,938 | 5,51E-16 | 3 | Dad1 |
| 3,53E-20 | 0,716949696 | 1 | 0,974 | 8,26E-16 | 3 | Nedd8 |

| | | | | | | |
|---|---|---|---|---|---|---|
| 5,60E-20 | 0,806678346 | 1 | 1 | 1,31E-15 | 3 | Rpl37a |
| 7,50E-20 | 1,375330379 | 1 | 0,566 | 1,76E-15 | 3 | Peg3 |
| 1,20E-19 | 0,699220521 | 1 | 0,761 | 2,82E-15 | 3 | Dnajc15 |
| 1,77E-19 | 1,068235194 | 1 | 1 | 4,14E-15 | 3 | Rps29 |
| 3,10E-19 | 0,678902034 | 1 | 0,985 | 7,26E-15 | 3 | Hmga1 |
| 5,89E-19 | 0,775449815 | 1 | 0,743 | 1,38E-14 | 3 | Lgals1 |
| 1,13E-18 | 0,684264675 | 1 | 0,996 | 2,65E-14 | 3 | Rpl28 |
| 4,46E-18 | 0,782118007 | 1 | 0,772 | 1,04E-13 | 3 | Nufip1 |
| 4,85E-18 | 1,038965886 | 1 | 1 | 1,14E-13 | 3 | Usmg5 |
| 7,33E-18 | 0,994066523 | 1 | 0,934 | 1,72E-13 | 3 | Sugt1 |
| 8,95E-18 | 1,093150833 | 1 | 0,993 | 2,10E-13 | 3 | Zfp985 |
| 1,36E-17 | 0,689631386 | 1 | 0,993 | 3,18E-13 | 3 | Sfr1 |
| 1,40E-17 | 0,736907656 | 1 | 1 | 3,27E-13 | 3 | Rpl18a |
| 2,56E-17 | 0,948378978 | 1 | 1 | 6,00E-13 | 3 | Rps28 |
| 3,54E-17 | 0,748746862 | 1 | 1 | 8,28E-13 | 3 | Sec61g |
| 4,03E-17 | 0,690261894 | 1 | 1 | 9,44E-13 | 3 | Gas5 |
| 6,62E-17 | 0,709033545 | 1 | 0,996 | 1,55E-12 | 3 | Nhp2 |
| 7,89E-17 | 0,771185097 | 1 | 0,919 | 1,85E-12 | 3 | Sparc |
| 8,25E-17 | 0,786509543 | 0,979 | 0,585 | 1,93E-12 | 3 | Nqo1 |
| 9,10E-17 | 0,863712137 | 1 | 1 | 2,13E-12 | 3 | Mif |
| 1,55E-16 | 0,91259944 | 1 | 1 | 3,63E-12 | 3 | Rpl36 |
| 1,79E-16 | 0,771037943 | 1 | 1 | 4,19E-12 | 3 | Rps15a |
| 2,28E-16 | 0,844809306 | 1 | 0,996 | 5,34E-12 | 3 | Gm13147 |
| 4,62E-16 | 0,737677065 | 1 | 0,949 | 1,08E-11 | 3 | Cyb5a |
| 5,56E-16 | 0,799629696 | 1 | 1 | 1,30E-11 | 3 | Uqcrq |
| 6,05E-16 | 0,816654487 | 1 | 1 | 1,42E-11 | 3 | Rps21 |
| 6,49E-16 | 0,70612518 | 1 | 1 | 1,52E-11 | 3 | Myl6 |
| 8,01E-16 | 0,870146435 | 1 | 0,967 | 1,88E-11 | 3 | Emb |
| 1,42E-15 | 0,690937466 | 1 | 1 | 3,32E-11 | 3 | Tmsb10 |
| 2,50E-15 | 0,832808539 | 1 | 1 | 5,86E-11 | 3 | Eif2s2 |
| 2,80E-15 | 0,979570338 | 1 | 0,824 | 6,55E-11 | 3 | Amd1 |
| 5,24E-15 | 0,835571985 | 1 | 0,993 | 1,23E-10 | 3 | Nap1l1 |
| 5,61E-15 | 0,892592317 | 0,957 | 0,57 | 1,31E-10 | 3 | Dnmt3a |
| 6,87E-15 | 0,750223098 | 1 | 0,989 | 1,61E-10 | 3 | Gm4204 |
| 9,75E-14 | 0,774255418 | 1 | 0,982 | 2,28E-09 | 3 | Gnl3 |
| 1,18E-13 | 1,024897471 | 1 | 0,996 | 2,77E-09 | 3 | Actg1 |
| 1,21E-13 | 0,961475981 | 1 | 0,96 | 2,84E-09 | 3 | Tmsb4x |
| 1,33E-13 | 0,684230243 | 1 | 0,996 | 3,11E-09 | 3 | Atp5g3 |
| 1,77E-13 | 0,9445126 | 1 | 0,923 | 4,15E-09 | 3 | Prdx6 |
| 2,38E-13 | 0,679279878 | 1 | 0,849 | 5,58E-09 | 3 | Sqstm1 |
| 2,59E-13 | 1,000442038 | 0,979 | 0,651 | 6,06E-09 | 3 | Gja1 |
| 6,28E-13 | 0,762372887 | 1 | 1 | 1,47E-08 | 3 | Rpl35 |
| 1,09E-12 | 0,684948937 | 1 | 0,982 | 2,55E-08 | 3 | Bex3 |
| 1,10E-12 | 0,750738625 | 1 | 0,96 | 2,58E-08 | 3 | Igf2bp1 |
| 1,37E-12 | 0,764847766 | 1 | 0,695 | 3,20E-08 | 3 | Tsc22d1 |
| 1,65E-10 | 0,704611407 | 1 | 0,864 | 3,86E-06 | 3 | Tex19.1 |
| 2,06E-10 | 0,67328379 | 1 | 1 | 4,83E-06 | 3 | Snrpg |
| 2,18E-10 | 0,854490115 | 1 | 1 | 5,10E-06 | 3 | Rpl5 |

| | | | | | | |
|---|---|---|---|---|---|---|
| 2,66E-10 | 0,804902504 | 1 | 1 | 6,23E-06 | 3 | Rpl39 |
| 4,93E-10 | 0,657453583 | 1 | 0,996 | 1,15E-05 | 3 | Zfp981 |
| 8,29E-10 | 0,943467511 | 1 | 0,985 | 1,94E-05 | 3 | Mt1 |
| 3,86E-08 | 0,661566019 | 1 | 0,989 | 0,000904962 | 3 | Pfdn1 |
| 2,90E-07 | 0,783621254 | 0,979 | 0,724 | 0,006781257 | 3 | Rhox5 |
| 3,14E-07 | 0,684735401 | 1 | 0,996 | 0,007346955 | 3 | AC121965.1 |
| 0,000308792 | 0,696074026 | 1 | 1 | 1 | 3 | Ndufa4 |

### Supplemental Theory

Fabrizio Olmeda and Steffen Rulands

December 30, 2020

In this Supplemental Theory we provide details of the calculations supporting the conclusions in the main text.

**6 Derivation of cross-correlation functions** **35**

**7 Stochastic simulations** **38**

### 1 Introduction

Novel technologies in single-cell genomics allow probing molecular states of several layers of cell fate regulation in single cells. Biological function, however, relies on collective processes in physical space and on the cellular scale. While measurements in genomics are restricted to the linear DNA sequence (sequence space), such mesoscopic processes rely on self-organisation in the *physical* space of the nucleus. In this Supplemental Theory we will we will give mathematical details on the methodology of how collective, mesoscopic processes in physical space can be inferred from measurements along the one-dimensional space of the DNA sequence.

To begin, we will define a general ansatz for the stochastic dynamics of the binding kinetics of interacting enzymes (Section 2). Constructing a coherent state path integral representation we will then infer the corresponding interaction kernel from the sequencing data. In Section 4 we will show how an effective description in physical space can be derived from the field theory in sequence space. In Section **??** we will finally use renormalisation group theory to derive correlations functions.

### 2 A general ansatz for enzyme-DNA kinetics

For simplicity in the following we will use the language of DNA methylation. The calculations we present here are, however, applicable to general enzyme-DNA kinetics. To infer the kinetics of de-novo methylation from the experimental sequencing data we begin with a general mathematical ansatz for the DNMT3 enzyme kinetics. In this ansatz, we first model the DNA sequence as a one-dimensional lattice. Each site on this lattice corresponds to a base pair. As only a fraction of sites are cytosines in a CpG context, and can therefore be methylated, we consider an equivalent but more efficient description only involving CpGs: in this description, each site of a one-dimensional lattice corresponds to a CpG and the heterogeneity in the distances between neighbouring CpGs is taken into account by appropriately chosen disordered parameters (see below). At each site, DNMT3 enzymes can bind to the DNA and, if bound, methylate this site. Although unbinding and potentially de-methylation processes may occur these processes do not influence any of the prediction we make and are therefore not explicitly stated simplicity in the following calculations. The emergence of a non-trivial power-law in average DNA methylation levels (Fig. 1 of the main text) and the power-law decay of the correlation function indicates that de-novo methylation is a collective process involving interactions of DNMT3 enzymes across extended genomic domains. We therefore make the ansatz that the binding rate of DNMT3 enzymes at a position $i$ is dependent on the presence of bound enzymes in its vicinity. Specifically, if the nearest bound sites are at positions $j_1$ and $j_2$ we write the binding rate as $J_{i,j_1} + J_{i,j_2}$, where the two terms correspond to the contribution from the left and right bound neighbour, respectively. Interaction with the closest occupied sites on the left and right, effectively restrict these interactions in range, cf. [2]. Omitting unbinding and demethylation terms for brevity the time evolution of the probability of finding a given DNMT3 binding and methylation profile, $P(\mathbf{D}, \mathbf{m}, t)$ follows a master equation of the form

$$\begin{aligned}\frac{\partial P(\mathbf{D},\mathbf{m},t)}{\partial t} =& \sum_{i=1}^{N}\sum_{l=i+1}^{N} J_{i,i+l}\left(\prod_{j=1}^{l-1}\bar{D}_{i+j}\right) D_l[D_i P(\bar{\mathbf{D}}_i,\mathbf{m},t) - \bar{D}_i P(\mathbf{D},\mathbf{m},t)] \\ &+ \text{l.n.n.} \\ &+ \text{demethylation and unbinding processes}\,,\end{aligned}$$

where $\mathbf{D}$ and $\mathbf{m}$ are a binary vectors describing DNMT3 occupancy and DNA methylation, respectively, such that, for example, $D_i = 1$ if site $i$ is occupied and $D_i = 0$ otherwise. $\bar{\mathbf{D}}_i$ is the same vector where, at position $i$, $D_i$ is replaced by $1 - D_i$ and

$\bar{D}_i = 1 - D_i$. $N$ is the number of lattice sites. The interaction with the left nearest neighbors has the same form as the term shown and is abbreviated by "l.n.n". In this ansatz, we implicitly assumed that there are no further non-linearities or spatial correlations relevant on large enough length scales, for example from the deposition of methyl groups by bound enzymes on the DNA or from de-methylation processes. We will show below that such additional processes are irrelevant under rescaling. In the context of de-novo methylation the non-linear nature of DNMT3 binding is supported by the literature [1].

In the following, as linear and uncorrelated processes do not contribute to the results obtained below, we will consider the marginal distribution

$$P(\mathbf{D}, t) = \sum_{m_i} P(\mathbf{D}|\mathbf{m}, t) P(\mathbf{m}, t) \,. \tag{1}$$

Following the definition of the model ansatz we have $P(\mathbf{D}|\mathbf{m}, t) = P(\mathbf{D}, t)$ such that the Master equation for the marginal distribution has the same form as above,

$$\frac{\partial P(\mathbf{D}, t)}{\partial t} = \sum_{i=1}^{N} \sum_{l=i+1}^{N} J_{i,i+l} \left( \prod_{j=1}^{l-1} \bar{D}_{i+j} \right) D_l [D_i P(\bar{\mathbf{D}}_i, t) - \bar{D}_i P(\mathbf{D}, t)] + \text{l.n.n.} \,. \tag{2}$$

As a side remark, it is interesting to note that instead of considering particle creation the model can be recast in terms of the creation of domains. In this picture, a binding event between two already occupied sites is equivalent to a fragmentation event splitting the domain into two smaller intervals. Restricting our attention to the the first moment, the we can rewrite our ansatz as a fragmentation equation,

$$\frac{\partial c(x, t)}{\partial t} = \int_0^\infty \mathrm{d}y F(x, y - x) c(y, t) - c(x, t) \int_0^x \mathrm{d}y F(y, x - y) \,, \tag{3}$$

where $c(x, t)$ is the number of domains of size $x$ at time $t$. A minimal fragmentation kernel for long range interactions the is $F(x,y) = 1/x^\lambda + 1/y^\lambda$. Defining the moments as $M_\alpha = \int \mathrm{d}x c(x,t) x^\alpha$ we obtain $M_{\alpha+\lambda+1} \sim t^{-(\alpha+\lambda)/(\lambda+1)}$, and the average occupancy, which is the zeroth moment, scales like $D \sim t^{1/(\lambda+1)}$. This equation describes the time evolution of the number of enzymes bound to the DNA. In the following we will give a rigorous derivation of this result as well as higher order moments, such as correlation functions, using field theoretical methods. These cannot be derived from simple mean field arguments.

### 3 Inference of de-novo methylation kinetics in sequence space

Having defined a general ansatz for the kinetics of emzyme binding we now infer the functional form of the interaction kernel from the sequencing data. To this end, we employ a coherent state path integral formulation of the master equation (1). Taking the semiclassical limit we will then compare the prediction for the first moment to the experimental data and we will analyze the effect of other possible processes that may contribute to the dynamics of de-novo DNA methylation such as processivity of DNMT3 enzymes. Although we here infer the interaction kernel from the time evolution of the first moment the methods works equally starting from the static measurements of the correlation function.

#### 3.1 Coherent state path integral formulation of the master equation

To begin, we construct a path integral representation of the master equation. This representation will allow us to compute first and higher order moments. Before proceeding with the path integral representation we introduce a Fock space, in which the probability distribution is formally written as,

$$|P(t)\rangle = \sum_{\mathbf{D}} P(\mathbf{D},t) a_1^{\dagger D_1} ... a_N^{\dagger D_N} |0\rangle \,. \tag{4}$$

The operator $a_i^{\dagger D_i}$ is the creation operator and formally represents a binding event at a given site. $D_i$ denotes the number of bound enzymes at site $i$. The creation and annihilation operators $a_i, a_i^\dagger$ act on the basis $|D\rangle$ as follows,

$$\begin{aligned} a_i^\dagger \,|D_i\rangle &= |D_i + 1\rangle \\ a_i \,|D_i\rangle &= D_i \,|D_i - 1\rangle \ , \end{aligned} \tag{5}$$

and they follow standard commutation rules $[a_i, a_i^\dagger] = 1$. Using this notation we can formally rewrite the master equation in terms of the creation operators, $a_i^\dagger$,

$$\partial_t |P(t)\rangle = -H|P(t)\rangle \,, \tag{6}$$

where $H$ is

$$H = -\sum_{i=1}^{N} \sum_{l=1}^{N-i} \prod_{j=1}^{l-1} \hat{\delta}_{D_{i+j,0}} \frac{\hat{\delta}_{D_{i+l,1}}}{l^\lambda} [a_i^\dagger \hat{\delta}_{D_{i,0}} - \hat{\delta}_{D_{i,0}}] + l.n.n.\,. \tag{7}$$

The terms $\delta_{D_{i,0}}$ are equal to 1 if an enzymes is not present at the CpG site $i$ and 0 otherwise and restrict binding to a single enzyme per site. With this, we can write the expectation value of any observable $A(\mathbf{D}, t)$ as

$$\langle A(\mathbf{D}, t)\rangle = \sum_{\mathbf{D}} A(\mathbf{D}) P(\mathbf{D}, t) \,. \tag{8}$$

We can rewrite this equation as

$$\langle A(\mathbf{D},t)\rangle = \langle 0|\prod_i e^{a_i} A(\mathbf{D})\,|P(t)\rangle\ . \tag{9}$$

where we introduce a coherent state basis $\langle 0|\,e^a$, which has the property to be the left eigenstate of the creation operator $a^\dagger$

$$\langle 0|\,e^a a^\dagger = \sum_{n=1}^{\infty} \frac{\langle 0|}{n!} a^n a^\dagger = \langle 0|\,e^a. \tag{10}$$

Within this basis, it is possible to write the field theory associated with the stochastic process after introducing the identity

$$1 = \int d\phi d\hat{\phi} e^{-\hat{\phi}\phi} e^{\phi a^\dagger}\,|0\rangle\,\langle 0|\,e^{\hat{\phi}a}, \tag{11}$$

which we will use later. In order to derive the field theoretical representation of the master equation we need to understand how the delta terms in $H$ act on the coherent state basis. Following the rules of [3], the $\hat{\delta}$ operator acts on the coherent state basis as

$$\begin{aligned} \langle\phi|\,a^\dagger\hat{\delta}_{\hat{n},m}\,|\phi\rangle &= \frac{1}{m!}\hat{\phi}\left(\hat{\phi}\phi\right)^m e^{-\phi\hat{\phi}}\,, \\ \langle\phi|\,a\hat{\delta}_{\hat{n},m}\,|\phi\rangle &= \frac{1}{(m-1)!}\phi\left(\hat{\phi}\phi\right)^{m-1} e^{-\phi\hat{\phi}}\,, \\ \langle\phi|\,\hat{\delta}_{\hat{n},m}\,|\phi\rangle &= \frac{1}{m!}\left(\hat{\phi}\phi\right)^m e^{-\phi\hat{\phi}}\,. \end{aligned} \tag{12}$$

In order to construct the field theory we first note that we can formally write a solution of the master equation as

$$|P(t)\rangle = e^{-Ht}|P(0)\rangle\,, \tag{13}$$

where $|P(0)\rangle$ is the initial state (i.e. the probability distribution of enzymes binding profiles at time $t = 0$). The exponential factor can be rewritten as

$$e^{-Ht} = (1 - \Delta t H)^{\frac{t}{\Delta t}} = (1 - \Delta t H) \cdot (1 - \Delta t H) \cdot \ldots . \tag{14}$$

By inserting the identity in the coherent state basis between every factor on the right hand side of Eq. (13), we find that the solution for any time $t_1$ can be written as

$$\begin{aligned} |P(t_1)\rangle = \int \prod_i d\phi_i(t_1 + \Delta t) d\hat{\phi}_i(t_1 + \Delta t) d\phi(t_1) d\hat{\phi}_i(t_1) e^{-\hat{\phi}_i(t_1)\phi(t_1)_i} \\ \cdot e^{-\hat{\phi}_i(t_1+\Delta t)\phi(t_1+\Delta t)_i \phi_i(t_1+\Delta t) a^\dagger} \\ \cdot |0\rangle \langle 0| e^{\hat{\phi}_i(t_1+\Delta t) a} (1 - \Delta t H) e^{\phi_i(t_1) a^\dagger} |0\rangle \langle 0| e^{\dot{a} t \phi_i(t_1) a} . \end{aligned} \tag{15}$$

In the previous equation we have to evaluate quantities in the coherent state basis between the bra and the ket,

$$\begin{aligned} \langle 0| e^{\hat{\phi}_i(t_1+\Delta t) a} (1 - \Delta t H) e^{\phi_i(t_1) a^\dagger} |0\rangle &= e^{\hat{\phi}_i(t_1+\Delta t)} e^{\phi_i(t_1)} - \Delta t \langle 0| e^{\hat{\phi}_i(t_1+\Delta t) a} (H) e^{\phi_i(t_1) a^\dagger} |0\rangle \\ &\approx e^{\hat{\phi}_i(t_1+\Delta t)} e^{\phi_i(t_1)} e^{-\Delta t H(\hat{\phi}_i(t_1), \phi_i(t_1))} , \end{aligned} \tag{16}$$

where $H(\hat{\phi}_i, \phi_i)$ is obtained by replacing all $a_i$ with $\phi_i$ and $a_i^\dagger$ with $\hat{\phi}_i$, using the previously introduced decomposition and by taking into account the rules for exclusion processes, Eq.(12). Repeating this procedure $t/\Delta t$ times for each factor $(1 - \Delta t H)$ we end up with an integral, $P_1$, over a product of three terms $P_2$, $P_3$, $P_4$. The integral reads

$$P_1 = \int \prod_i d\hat{\phi}_i(t) d\phi_i(t) d\hat{\phi}_i(t - \Delta t) d\phi_i(t - \Delta t) \cdot \ldots \cdot d\hat{\phi}_i(\Delta t) d\phi_i(\Delta t) d\hat{\phi}_i(0) d\phi_i(0) \ldots , \tag{17}$$

which can be rewritten compactly as a functional integral

$$P_1 = \int \mathcal{D}[\phi]\mathcal{D}[\hat{\phi}] \ldots . \tag{18}$$

$P_2$ is composed of a product of terms which can be rewritten by means of Riemann integration,

$$P_2 = \prod_{t_1=\Delta t}^{t} e^{\hat{\phi}(t_1+\Delta t)\phi(t_1)-\hat{\phi}(t_1)\phi(t_1)} \approx e^{-\int dt \partial_t \hat{\phi}\phi}. \tag{19}$$

Finally there are further $t/\Delta t$ terms coming from the Hamiltonian evaluated at each time step,

$$P_3 = \prod_{t_1=\Delta t}^{t} e^{-\Delta t H(\hat{\phi}(t_1),\phi(t_1))} \approx e^{-\int dt H(\hat{\phi}(t),\phi(t))} . \tag{20}$$

The final factor, $P_4$, represents initial conditions and we refer to Ref. [6] for a discussion of this term. With this, any observable can then be expressed as a path integral of the form

$$A(\mathbf{D}) = \int \mathcal{D}[\phi]\mathcal{D}[\hat{\phi}] A(\boldsymbol{\phi}, \hat{\boldsymbol{\phi}} = 1) e^{-S[\hat{\phi},\phi]} , \tag{21}$$

with

$$S[\hat{\phi}, \phi] = -\sum_i \phi_i(t_f) + \int_0^{t_f} \mathrm{d}t \sum_i \left( \hat{\phi}_i(t)\partial_t \phi_i(t) + H_i[\hat{\phi}, \phi] \right) , \tag{22}$$

where we have performed a partial integration in time. The Hamiltonian in this action reads

$$H_i[\hat{\phi}, \phi] = (1 - \hat{\phi}_i) e^{-\hat{\phi}_i \phi_i} \left[ \sum_{l=1}^{N-i} \frac{\hat{\phi}_{i+l}\phi_{i+l}}{l^\lambda} e^{-\sum_j \hat{\phi}_{i+j}\phi_{i+j}} + \sum_{l=1}^{i} \frac{\hat{\phi}_{i-l}\phi_{i-l}}{l^\lambda} e^{-\sum_j \hat{\phi}_{i-j}\phi_{i-j}} \right] . \tag{23}$$

Defining the generating functional of correlations, $Z[\mathbf{h},\phi,\bar{\phi}]$, as

$$Z[\mathbf{h},\phi,\bar{\phi}] = \int \mathcal{D}[\mathbf{h},\phi,\bar{\phi}] e^{-S[\phi,\bar{\phi}]+\int_0^y ds \int_0^{t_f} dt \left[h(s,t)\phi(s,t)+\bar{h}(s,t)\bar{\phi}(s,t)\right]}\,, \tag{24}$$

expectation values of products of observables, such as correlation functions, can then be expressed as functional derivatives with respect to the auxiliary external field,

$$\langle \phi(s,t)\phi(y,t')\rangle = \frac{\delta^2}{\delta h(s,t)\delta h(y,t)} Z[\mathbf{h},\phi,\bar{\phi}]|_{\mathbf{h}=0}\,. \tag{25}$$

#### 3.2 Semiclassical limit of the field theory

In this section we infer the functional form of the interaction kernel of enzyme binding events from a semiclassical solution of the field theory. In a first step, we rewrite the Hamiltonian (23) in continuous space. This is the same as we have done previously with Riemann integration. The idea is that by introducing a spatial discretisation $\sum_i \Delta s \to \int ds$ the Hamiltonian in the action, Eq.(23), becomes

$$\begin{aligned} H[\hat{\phi},\phi] = (1-\hat{\phi}(s))e^{-\phi\hat{\phi}}\Big[ \int_0^{N-s} \mathrm{d}y \frac{\hat{\phi}(s+y)\phi(s+y)}{y^\lambda} e^{-\int_{z=0}^{y}\mathrm{d}z \hat{\phi}(s+z)\phi(s+z)} \\ + \int_0^s \mathrm{d}y \frac{\hat{\phi}(s-y)\phi(s-y)}{y^\lambda} e^{-\int_{z=0}^{y}\mathrm{d}z\hat{\phi}(s-z)\phi(s-z)}\Big]\,. \end{aligned} \tag{26}$$

Expanding to first order in the exponential, to the second order in the terms $\hat{\phi}(s\pm y)\phi(s\pm y)\Big/y^\lambda$ and extending the limit of integration to infinity the Hamiltonian obtains the form

$$H[\hat{\phi},\phi] = J\Gamma(1-\lambda)\left(1-\hat{\phi}\right)e^{-\phi\hat{\phi}}\left[2\left(\hat{\phi}\phi\right)^\lambda + \left(\phi\hat{\phi}\right)^{\lambda-3}(2-3\lambda+\lambda^2)\frac{\partial^2\left(\phi\hat{\phi}\right)}{\partial s^2}\right]\,. \tag{27}$$

By minimising the action, $\delta S\Big/\delta\phi(\hat{x})\,|_{\hat{\phi}(x)=1} = 0$ while setting $\hat{\phi}(x)=1$ for probability conservation we obtain a partial differential equation describing the time evolution of the DNMT3 binding profile, $\phi(x)$, in the semi-classical limit,

$$\frac{\partial \phi(s)}{\partial \tilde{t}} = \phi(s)^{\lambda} + D\phi(s)^{\lambda-3}\partial_s^2\phi(s) + \eta(s,t)\,. \tag{28}$$

Here, $\tilde{t} = 2Jt\Gamma(1-\lambda)$ and $D = (2-3\lambda+\lambda^2)/2 > 0$. In the hard-boson path-integral representation, Eq. (26), the term $\exp\left[-\int_{z=0}^{y} \mathrm{d}z\hat{\phi}(s+z)\phi(s+z)\right]$ in the Hamiltonian, for a slowly varying field can be approximated by $e^{-y\langle\hat{\phi}(s)\phi(s)\rangle}$, where $\langle\hat{\phi}(x)\phi(x)\rangle$ is the average product of the fields. This defines an effective exponential cutoff to the interactions at a characteristic length $1/y^{\lambda}$. This interpretation of the exponential term justifies the expansion within the limit of integration.

#### 3.3 Inference of the interaction kernel

To infer the interaction kernel $J_{i,i+l}$ we compute first and higher order moments of the local methylation density $m(s,t)$ which, in the semiclassical limit, follows $\partial_t m(s,t) = k\phi(s,t)$. Having computed these moments for a general class of interaction kernels we can then match theoretical predictions with experimental data to infer the functional form of interactions between enzyme binding events. Here, we focus on the time evolution first moment of the global DNA methylation level, $m(t) = 1/N\sum_{s=1}^{N} m(s,t)$. We will later use the prediction of higher order moments to validate our results. An equally feasible approach would be to infer the interaction kernel from higher moments, such as the correlation functions in Eq. (76), and use predictions of the time evolution of the first moment to verify results. To begin, we sum Eq. (28) to obtain a differential equation for $\phi(t) = 1/N\sum_{s=1}^{N}\phi(s,t)$, which again is solved by

$$\phi(t) = t^{1/(1-\lambda)}\,. \tag{29}$$

Taken together, average DNA methylation increases according to

$$m(t) = m(t=0) + k\frac{1-\lambda}{2-\lambda}t^{1+1/(1-\lambda)}\,. \tag{30}$$

Therefore, in order to match the experimentally obtained exponent of 5/2 we find that $\lambda = 1/3$, such that the interaction kernel reads

$$J_{i,i+l} = \frac{1}{l^{1/3}}\,. \tag{31}$$

This means, that the total binding rate at a position $i$ is given by $1/l_L^{1/3} + 1/l_R^{1/3}$, where $l_L$ and $l_R$ are the distances to the left and right nearest bound enzymes, respectively. The total binding rate in a region of size $l$ is then given by the sum $\sum_{s=1}^{l-1}\left[1/s^{1/3} + 1/(l-s)^{1/3}\right]$, which scales like $l^{2/3}$.

#### 3.4 Irrelevance of processive DNA methylation on large spatial scales

As shown in Ref. [4] in a simpler version of this model, there exist a continuous phase transition to the absorbing state for vanishing values of the enzyme unbinding rate, $u$, and for $\lambda > 1$. In our case, where $\lambda = 1/3$, classical renormalisation group arguments cannot be applied. We can, however, still investigate fluctuations around a steady state solution given by a balance between binding and unbinding events, a mathematical assumption that is needed to avoid a trivial solution where all sites or occupied. A term descibing the unbinding of enzymes will be of the form $H_u = u\sum_{i=1}^{N}(1-a_i)\hat{\delta}_{D_i,1}$ which results in a term in the action of the form $u\phi(\hat{\phi}-1)e^{-\hat{\phi}\phi}$. Starting with Eq. (27), dimensional analysis yields in the momentum $[k]$ and time scale $[\tau]$,

$$[\hat{\phi}] = [k]^0[\tau]^0, [\phi] = [k]^d[\tau]^0[\tau]^0, [S] = [k]^0[\tau]^0, [u] = [k]^0[\tau]^{-1}. \tag{32}$$

Note that we choose $\hat{\phi}$ to be adimensional. Moreover the term which accounts for exclusion, $e^{-\phi\hat{\phi}}$, whose exponent must be adimensional, has been implicitly written as $e^{-v\phi\hat{\phi}}$ following [5], where $v = [k]^{-1}$ is the lattice discretization. In spatial dimension $d$ we obtain from Eq. (27) a system of equations

$$[k]^{d(\lambda-3)}[k]^2[J][\tau] = [k]^0[\tau]^0\,, [k]^{-d}[k]^{d\lambda}[J][\tau] = [k]^0[\tau]^0\,, \tag{33}$$

with solutions

$$[J] = [\tau]^{-1}[k]^{-d(\lambda-3)+2}\,, [J] = [\tau]^{-1}[k]^{d(1-\lambda)}\,, \tag{34}$$

which are self-consistent only for $d = 1$ and $[J] = [k]^{1-\lambda}[\tau]^{-1}$. This scaling comes from the fact that the diffusion constant is not an independent parameter but depends on the local binding rate. We will now show that processivity of DNMT3 enzymes becomes irrelevant under renormalisation. The term representing processivity in the action is of the form $D_0\hat{\phi}\partial_s^2\phi$, with scaling dimensions $[D_0] = [k]^{-2}[\tau]^{-1}$. This term must be compared to the effective diffusion $[J] = [k]^{1-\lambda}[\tau]^{-1}$. Processivity is then irrelevant for $\lambda < 3$, which is satisfied in our case. Even though this argument is valid in scenarios where binding and unbinding of DNMT3 enzymes are roughly balenced (i.e. close to he phase transition), we do not expect processivity to play a dominant role in the scaling and leading order of correlation functions and propagators away from the phase transition.

#### 4 From sequence space to physical space

Having defined the kinetics of de-novo methylation in sequencing space we will now derive an effective theory for the collective dynamics in physical space. As in our model DNA methylation events do not introduce additional spatial correlations, cf. Section 2, we will in this section use the terms DNMT3 binding profile and DNA methylation profile interchangeably.

#### 4.1 Derivation of the field theory in physical space

Before rigorously deriving the effective time evolution in physical space in technical terms we first give a heuristic explanation for the structure of the resulting partial differential equation.

##### 4.1.1 A heuristic motivation

Starting from the Langevin equation (28) in the limit of low noise, we make an ansatz to study the flux of methylated CpGs in physical space, denoted by the coordinate $x$. To begin, we rewrite (28) in terms of a field $h(x,t)$ defined as the perturbation around a global homogeneous average, $\phi_0(t)$, $\phi(x,t) = \phi_0(t) + h(x,t)$, such that the corresponding Langevin equation reads to second order in $h(s,t)$

$$\frac{\partial h(s,t)}{\partial t} = bh(s,t) + c^- h(s,t)^2 + D_0 \partial_s^2 h(s,t) + D_1^- h(s,t) \partial_s^2 h(s,t)\,. \tag{35}$$

$b$, $c^-$, $D_0$, and $D_1^-$ are constant terms coming from the Taylor expansion of Eq. (28) with $b = \lambda(\phi_0)^{\lambda-1}$, $c^- = \phi_0^{\lambda-2}\lambda(\lambda-1)/2$, $D_0 = D\phi_0^{\lambda-3}$, and $D_1^- = D(\lambda-3)\phi_0^{\lambda-4}$.

In physical space, changes in the topology of the DNA due to de-novo methylation kinetics lead to an additional flux term, $J$, that must conserve total DNA methylation,

$$\frac{\partial h}{\partial t} = bh(x,t) + c^- h(x,t)^2 + D_0 \partial_x^2 h(x,t) + D_1^- h(x,t) \partial_x^2 h(x,t) - \partial_x J[h(x,t), \partial_x h(x,t), \dots]\,. \tag{36}$$

This flux term can only depend on powers of $h(x,t)$ and its spatial derivatives. Algebraic terms in $h(x,t)$ correspond to active transport of methylated sites in the nucleus, which is not biologically reasonable. To lowest order, the flux term therefore takes the

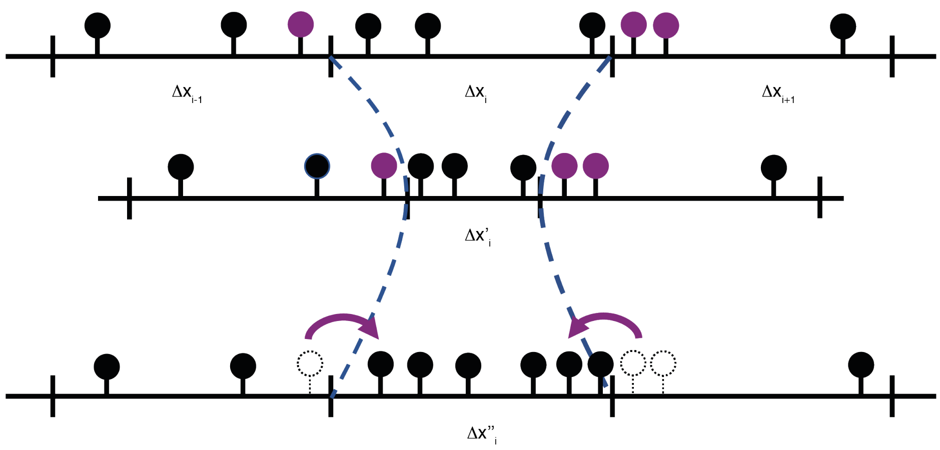

Figure 1: Effect of changes in DNA compaction on the redistribution of methylated CpGs in physical space (black and purple circles). The method consists of two steps: first, after a methylation event the chromatin gets contracted (blue dashed line). We then renormalise space such that the total length of the system remain invariant (blue dashed line). With this procedure methylated sites of neighboring domains (purple circles) effectively lead to a flux into the contracted domain in physical space.

form

$$J = D_c \partial_x h(x,t) + \xi(x,t)\,, \tag{37}$$

where $\xi$ is uncorrelated Gaussian white noise with $\langle \xi \rangle = 0$ and correlation $\langle \xi(x,t)\xi(x',t') \rangle = 2\Gamma\delta(t-t')\delta(x-x')$. Taken together we find

$$\frac{\partial h}{\partial t} = bh(x,t) + c^{-}h(x,t)^2 + (D_0 - D_c)\partial^2 h(x,t) + D_1^{-} h(x,t)\partial_x^2 h(x,t) - \partial_x \xi(x,t)\,. \tag{38}$$

Therefore, according to this heuristic argument, the dynamics in physical space is described by the Edward-Wilkinson (EW) equation with additional non-linear terms which are irrelevant under renormalization . In the following we will derive rigorously the flux terms and the noise as well as the coefficient $D_c$.

##### 4.1.2 Rigorous derivation of the dynamics in physical space

To systematically derive the dynamics in physical space from the inferred kinetics in sequence space we start with the partial differential equation describing the spatio-temporal dynamics in sequence space in the semiclassical limit, Eq. (28). To begin, we will ask how small length elements in physical space evolve in time for a given position in sequence space. Based on this, we will calculate effective local fluxes in DNA methylation density in physical space. Specifically, we seek to define a function $g_i$ that describes the evolution of length elements in a properly defined physical space with respect to changes in DNA methylation density,

$$\delta\Delta x_i = \Delta x_i - g_i(\Delta x_i), \tag{39}$$

with initial conditions $\Delta x^0 = \Delta s$. $\Delta s$ and $\Delta x$ are, respectively, length elements in sequence and physical space.

Starting from a description in sequence space, given by the semiclassical time evolution of the field $\phi(s,t)$, we aim to derive a master equation in physical space. Let $\Delta\xi_i$ be the length of a discrete element in physical space. A DNA methylation event (understood in a coarse-grained fashion) causes a local compaction of the DNA, such that this length element is contracted by

$$\Delta\xi_i' = \Delta\xi_i^{1/3}\,. \tag{40}$$

In the absence of demethylation continous de-novo methylation will therefore continously compact the DNA locally such that the total length of the DNA in physical space, $\Lambda = \sum_i \Delta\xi_i$ decreases over time. This gives rise to a flux with a velocity that locally depends on the entire concentration field left and right of a given position. To avoid such difficulties, we define a dynamic renormalisation scheme with renormalised length elements $\Delta x_i$ such that the total length of the DNA, $L = \sum_i \Delta x_i$, remains constant over time. To achieve this, for a given de-novo methylation event, $\Delta x_i$ is first contracted according to $\Delta x_i' = \Delta x_i^{1/3}$ and then rescaled by $\Delta x_i'' = b\Delta x_i'$, with the rescaling factor $b > 1$ given by

$$b = \frac{\Delta x_i + \Delta x_{i+1} + \Delta x_{i-1}}{\Delta x_i' + \Delta x_{i+1}' + \Delta x_{i-1}'} \,. \tag{41}$$

After renormalizing back such that the total length is unchanged, we get an effective flux of methylated sites (Fig. 1), close to the contracted domain. With this, we obtain an updating scheme for the concentration of methylated sites at position $i$, $\rho_i$, at each DNA methylation event. Specifically, the updated concentration at a given position, $\rho_i'$ is given by contributions from the original concentrations and symmetric fluxes from the adjacent left and right length elements,

$$\rho_i'' = \rho_i + (\rho_{i+1} + \rho_{i-1}) \frac{\Delta x - b\Delta x^{1/3}}{2\Delta x} \,. \tag{42}$$

We now consider the joint probability $P(\boldsymbol{\rho}, t)$ to find a given concentration profile $\boldsymbol{\rho}$ at a time $t$. On time scales much larger than the time scales associated with microscopic DNA methylation and compaction events, we can define the rate of de-novo methylation in a given length element $i$, $W(\rho_i) = \rho_i^\lambda$ with $\lambda = 1/3$. In this limit, the time evolution of $P(\boldsymbol{\rho}, t)$ is then given by a master equation for the redistribution of DNA methylation marks in physical space. It takes the form

$$\begin{aligned} \partial_t P(\boldsymbol{\rho}) = &\sum_i \left[W(\rho_i - r\rho_{i-1}) P(\rho_i - r\rho_{i-1}, \rho_{i-1} + r\rho_{i-1}, \boldsymbol{\rho})\right] \\ &+ \sum_i \left[W(\rho_i - r\rho_{i+1}) P(\rho_i - r\rho_{i+1}, \rho_{i+1} + r\rho_{i+1}, \boldsymbol{\rho})\right] \\ &- 2\sum_i W(\rho_i) P(\boldsymbol{\rho}, ) \,, \end{aligned} \tag{43}$$

where $r = (\Delta x - b\Delta x^{1/3})\big/(2\Delta x)$ is a dimensionless parameter describing the effective flux of DNA methylation in physical space due to a methylation event. In this notation, $P(\boldsymbol{\rho})$ is the probability of a given density profile, $P(\rho_{i+1} - 1, \boldsymbol{\rho})$ stands for the probability of that density profile $\boldsymbol{\rho}$, where at the position $i+1$, the density is $\rho_{i+1}-1$. The term $W(\rho_i)$ accounts for the rate of the DNA methylation event $W(\rho_i) = \rho_i^\lambda$ (i.e. the binding rate). This master equation can be rewritten in terms of lowering and raising operators as

$$\partial_t P(\boldsymbol{\rho}) = \sum_i \left[L_i^{-r\rho_{i-1}} L_{i-1}^{r\rho_{i-1}} + L_i^{-r\rho_{i+1}} L_{i+1}^{r\rho_{i+1}} - 2\right] W(\rho_i) P(\boldsymbol{\rho}) \,, \tag{44}$$

where a general operator $L^{\pm\rho_i}$ acts on functions to its right as $L^{\pm\rho_i} f(m_i) = f(m_i \pm \rho_i)$. This operator can be identically rewritten as $L_i^{-r\rho_{i-1}} = e^{-r\rho_{i-1}\partial_{\rho_i}}$ and expanded as $L_i^{-r\rho_{i-1}} = 1 - r\rho_{i-1}\partial_{\rho_i} + \mathcal{O}(r^2)$. We now proceed with a linear noise approximation of the Master equation, where the observable $\rho_i$ is split into a deterministic and a stochastic part

$$\rho_i = N\phi_i + \sqrt{N}\eta_i \,. \tag{45}$$

$N$ is the system size. Written in this form the operators become $L_i^{-r\rho_{i-1}} = 1 - rN^{-1/2}\rho_{i-1}\partial_{\eta_i} + \mathcal{O}(N^{-1})$. The terms on the right hand side of the Master equation are then to lowest order in $N$, $\mathcal{O}(1/\sqrt{N})$,

$$\sum_i r\left[\phi_{i-1}(-\partial_{\eta_i} + \partial_{\eta_{i-1}}) + \phi_{i+1}(-\partial_{\eta_i} + \partial_{\eta_{i+1}})\right] W(\phi)\Pi(\eta) \,. \tag{46}$$

As fluctuations in $\rho$ are given by fluctuations in $\eta$ we have $dP(\rho) = d\Pi(\eta)$, meaning that the probability distribution of the entire process is solely determined by its stochastic part. We now take a continuum approximation such that $\partial_{\xi_{i+1}} \approx \partial_{\eta(x)} \pm a_0\partial_x\partial_{\eta(x)}$ and the same for $\phi_{i+1}$, where $a_0$ is the lattice spacing, i.e. the distance between two base pairs. After these steps we obtain for the right hand side of the Master equation,

$$\int dx r a_0^3 \partial_x \phi(x) \partial_x \left[ W(\phi(x)) \frac{\delta \Pi(\eta)}{\delta \eta} \right] \ . \tag{47}$$

By applying the same procedures to the left hand side of the Master equation to the same order in $N$ we obtain

$$\partial_t P(\rho) = \partial_t \Pi - \sqrt{N} a_0 \int dx \, \frac{d\phi(x)}{dt} \frac{\delta \Pi}{\delta \eta} \, , \tag{48}$$

and by integrating by parts the right hand side and taking equal orders on both side of the expanded master equation, we obtain a change in the density field $\phi(x,t)$ of the form

$$\partial_t \phi(x) = -r a_0^2 W(\phi(x)) \partial_x^2 \phi(x) \, . \tag{49}$$

This partial differential equation describes the flux of DNA methylation density in physical space stemming from the local, methylation-dependent compaction of the DNA. How do the other terms in (28) evolve in physical space? In general, the procedure outlined above leads to a change in the functional form of long-range interactions in physical space. On the mean-field level, however, such interactions again give rise to a local and a diffusion term with potentially different non-linear dependencies on $\phi(x,t)$. Following the calculations we performed in sequence space in Section 2 the exponents describing these non linearities in the Langevin equation are not independent of each other. As the first moment as a global average must be identical in sequence space and in physical space the local term, and therefore also the diffusive term, must have the same form in sequence and in physical space. Therefore, in physical space, the time evolution of $\phi(x,t)$ is described by a combination of processes identical in sequence space and an additional term, Eq. (49), describing the flux of DNA methylation in physical space due to changes in DNA topology. Taken together, after substituting the definition of $W(\phi)$, $W(\phi(x)) = \phi(x)^\lambda$, we arrive at a partial differential equation describing the time evolution of $\phi(x,t)$ physical space,

$$\partial_t \phi(x,t) = \phi(x,t)^\lambda + \phi(x,t)^{\lambda-3}\partial_x^2\phi(x,t) - r\phi(x,t)^\lambda \partial_x^2 \phi(x,t)\,. \tag{50}$$

By taking into account the next highest order in the Van Kampen expansion we derive a term for the noise, which due to the conservation of methylation in the renormalisation procedure is entirely conservative,

$$\partial_t \phi(x,t) = \phi(x,t)^\lambda + \phi(x,t)^{\lambda-3}\partial_x^2\phi(x,t) - r\phi(x,t)^\lambda \partial_x^2 \phi + \eta(x,t) + \partial_x\left[g(\phi(x,t))\xi(x,t)\right] + \ldots\,. \tag{51}$$

The noise terms have correlations $\langle \xi(x,t)\xi(x',t')\rangle = 2\Gamma_C \delta(t-t')\delta(x-x')$ and $\langle \eta(x,t)\eta(x',t')\rangle = 2\Gamma_{NC} f(\phi(x,t))\delta(t-t')\delta(x-x')$, where the latter comes from terms proportional to $\hat{\phi}(x,t)^2$ in the field theoretical description (26). As we will be in the following interested in perturbations around a dynamical homogeneous solutions the particular dependencies in $g(\phi)$ and $f(\phi)$ will not be relevant for the remainder of our analysis.

The partial differential equation describing the time evolution of the field $\phi(x,t)$ is structurally similar to Eq. (28), but contains a new term $-r\phi(x,t)^\lambda \partial_x^2 \phi$ which is an anti-diffusive term counteracting the diffusion term. We expect that with increasing values of $\phi$ this term will dominate the diffusion term and potentially lead to the formation of highly methylated regions in physical space. Although we did not explicitly state higher order terms in the spatial derivatives these terms must exist and in the next section we will see how such terms affect the formation of methylation condensates. Considering perturbation of the previous equation $\phi(x,t) = \phi_0 + h(x,t)$ we arrive to the equation we intuitively derived before Eq. (38).

#### 4.2 Formation of condensates in physical space

Eq. (51) highlights how the dynamics of methylated domains in physical space may let new physics emerge with respect to the dynamics in sequence space. In this section, we will systematically investigate whether spatial structures emerge in physical space. To this end, we will investigate whether a homogeneous field in physical space is linearly unstable, which would imply the emergence of a characteristic length scale resembling DNA methylation condensates. To this end we take into account the next highest order term in $\phi$, $\epsilon\partial_x^4\phi$, which describe restoring forces counteracting DNA compaction at a finite length scale. Condensation happens if a spatial perturbation, $\delta\phi$, of a homogeneous solution, $\phi_0$, is unstable. Following standard procedures [14] we linearised Eq. (50) and made a general ansatz for the time evolution of the field $\phi(x,t)$ after perturbation with wave vector $k$,

$$\phi(x,t) = \phi_0 + e^{\omega t}e^{ikx}\delta\phi \tag{52}$$

The homogeneous state is unstable if $\omega > 0$. We obtain a general dispersion relation, which relates the rate of growth of the instability to the wavelength of the perturbation, of the form

$$\omega(k) = \lambda\phi_0^{\lambda-1} - \left(\phi_0^{\lambda-3} - r\phi_0^{\lambda}\right)k^2 - \epsilon^2 k^4\,. \tag{53}$$

The condition for formation of clusters of methylated DNA is then that the maximum of this function is greater than zero for some values of $k$ and its second derivative is negative, such that this maximum occurs at a finite value of $k$ and $\omega(k=0) > 0$. We obtain an instability if $\phi_0 > r^{\frac{1}{\lambda-3}}$. In Fig(2), we show the dispersion relation having fixed $r$ with varying $\phi_0$. The strongest growing mode at the point of the instability is $k = \frac{\sqrt{r^{7/8}-r}}{\sqrt{2\epsilon}}$, which gives an indication of the expected typical length scale of the resulting pattern. In summary, we expect an instability in the form of finite size

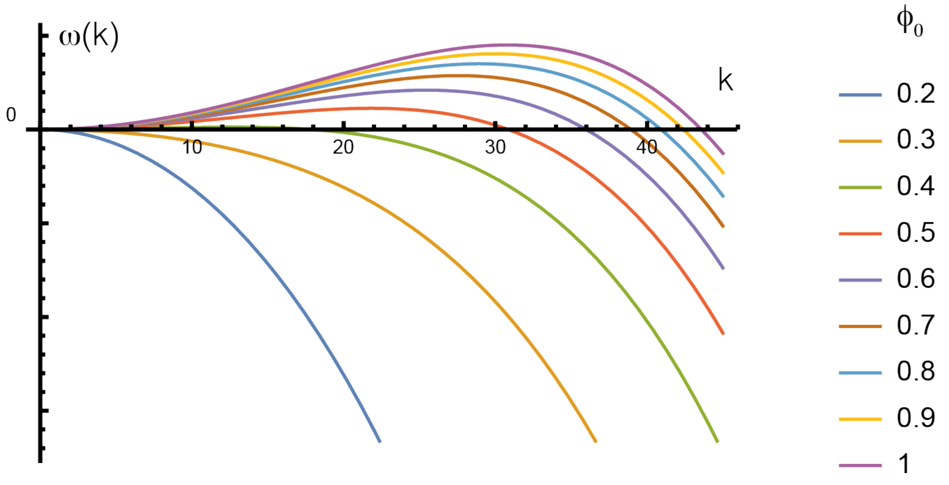

Figure 2: Dispersion relation of Eq.(51) with added a fourth order term in the derivative $\epsilon\partial_x^4\phi(x,t)$ and parameters value $r = 20, \epsilon = 0.01$

methylation condensates to arise if the average DNA methylation concentration, $\phi_0$, exceeds a threshold given by $r^{\frac{1}{\lambda-3}}$.

#### 4.3 Order of magnitude estimate of condensate sizes

As the precise values of $r$ and $\epsilon$ are unknown the linear stability analysis cannot be used straight forwardly to identify the length scale of the predicted methylation condensates. To get an order of magnitude estimate of these condensates we therefore resort to dimensional analysis. There are three length scales involved in the formation of methylation condensates corresponding to the parameters determining the dispersion relation in the previous section:

1. The typical distance between methylated CpGs, $l_{5mC}$. With an average CpG density of roughly 1% and average DNAme level of 50% we estimate that $l_{5mC} \approx$ 500 bp.

2. The typical length scale over which DNAme locally affects DNA compaction, $l_c$. From the cross-correlation function between DNAme and accessibility we find

that $l_c \approx 1000\,\mathrm{bp}$, cf. Fig. S3B. We expect that this length scale affects the size of condensates positively.

3. A length scale describing the restoring force counteracting DNA compaction, $l_r$. We expect this to be of the same order of magnitude as the DNA persistence length, $l_r \approx 100\,\mathrm{bp}$, and to affect the size of condensates negatively.

Taken together, in order to create another length scale reflecting the typical size of condensates from these three length scales, we find $l \approx l_{5mC} l_c / l_r$ which is approximately equal to 5000bp.

### 5 Derivation of the correlation function

In this section, we give details of the calculation of the correlation function. To illustrate the idea behind the renormalisation group procedure we will first, for didactic reasons, calculate the correlation function of a simplified model in physical space, Eq. (38). In a second step, we will then turn to the non-local dynamics in sequence space. The (connected) correlation function for the enzyme binding profile reads, $\langle \phi(x,t)\phi(x',t)\rangle - \langle \phi(x,t)\rangle\langle \phi(x',t)\rangle$, and, as the step involving the actual chemical modification of the DNA does not introduce further spatial correlations, is equal to the correlation function of the DNA methylation profile.

#### 5.1 An instructive example in physical space

To compute scaling exponents of the correlation function, we begin by performing renormalization of Eq.(38). The idea of the renormalisation group in momentum space is as follows: First, we transform the equation to be renormalised to Fourier space,

$$h(k,\omega) = G_0(k,\omega)\left[\xi(k,\omega) + \int_{\omega'} form, t_q W(k,q) h(q,\omega) h(k-q), \omega - \omega')\right], \quad (54)$$

with $h(k,\omega) = \int d\mathbf{x} \int dt h(x,t)\, e^{ikx} e^{i\omega t}$, vertex $W(k,q) = D_1^-(k-q)^2$, propagator $G_0(k) = \left(i\omega + \tilde{D}_0 k^2\right)^{-1}$ and $\tilde{D}_0 = D_0 - D_c$. The integral over momenta $k$ must be than taken in $(0,\Lambda)$ (Brillouine zone), where $\Lambda$ is the inverse of the lattice spacing (1 base pair). First, we divide the interval $k \in (0,\Lambda)$ into two different intervals, $k \in (0, \Lambda e^{-l}))$ and $k \in (\Lambda e^{-l}, \Lambda)$, with $l \ll 1$. This is done, because integrals coming from the self consistency equation (54) are often divergent as $k \to 0$ below the critical dimension. In the following we will discuss Wilson's momentum shell approach to deal with this problem of infrared divergences. The first step is to integrate out short wavelength (high momenta) on the momentum shell $k \in (\Lambda e^{-l}, \Lambda)$ which leaves us with an integral over $(0, \Lambda e^{-l})$. In a second step we rescale momenta by $k \to k e^{-l}$ to make length scales comparable, which again implies rescaling of other dimensional quantities in order to leave Eq. (38) invariant. Defining $b = e^l$ the rescaling step gives

$$t = b^z t', \; x = bx', \; \rho = b^\chi \rho' \,. \quad (55)$$

In order to perform the integration on the momentum shell we perform standard renormalisation group procedures [8] by perturbatively solving Eq. (54) for $h(x,t)$ and quantities derived from it, such as the propagator, correlator and vertex. These perturbative solutions can be represented in terms of Feynman diagrams as in Fig. 3. Every vertex represent an integral over momenta, the sum of which must be equal to zero. After evaluating these diagrams we get the following flows for the parameters (other parameters become irrelevant for the exponents under renormalisation, e.g.

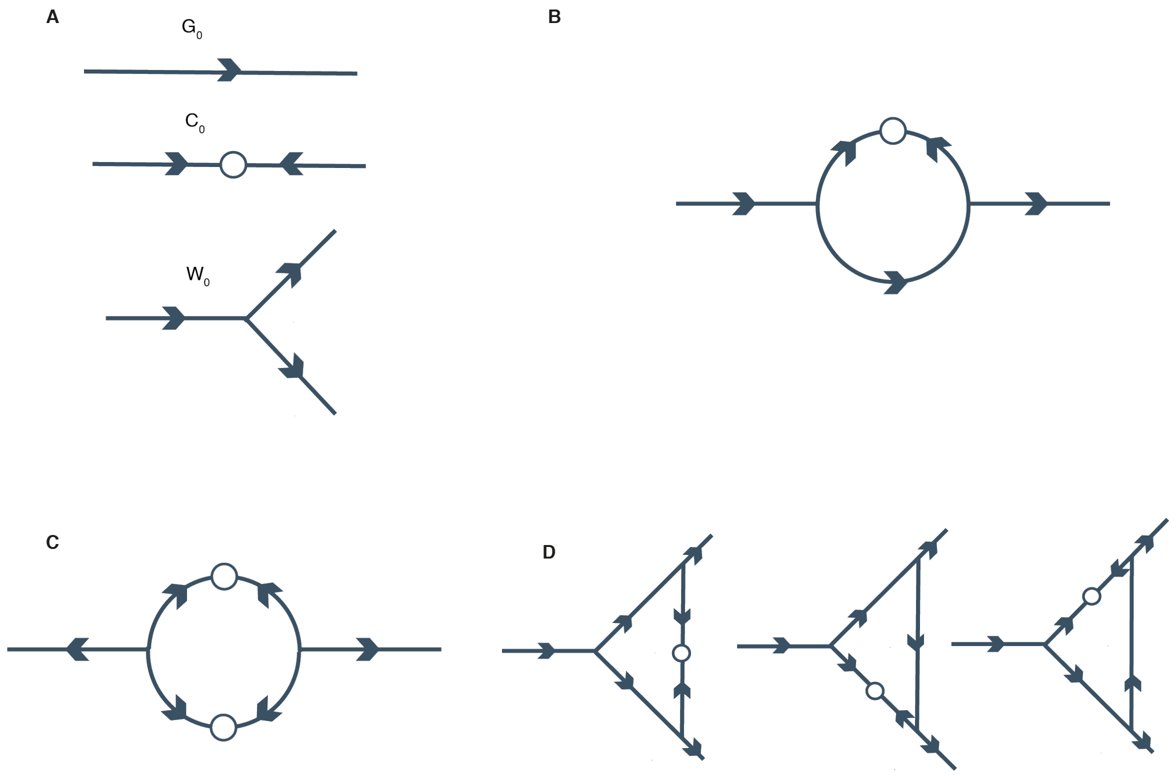

Figure 3: (A): Bare propagator ($G_0$), correlator ($C_0$) and vertex ($W_0$). ($B, C, D$): one loop correction to the propagator, correlator and vertex

terms in higher order spatial derivatives),

$$\begin{aligned} \partial_l \tilde{D}_0 &= [z-2]\, D_0 \\ \partial_l D_1^- &= [z+\chi-2]\, D_1^- \\ \partial_l \Gamma_C &= [z-2\chi-d-2]\, \Gamma_C\,, \end{aligned} \tag{56}$$

All quantities are computed in the limit of long time scales, $\omega \to 0$, and we will drop the explicit $\omega$ dependence in the following.

In the hydrodynamic limit, $k \to 0$ all integrals over the momentum shell evaluate to 0 leading to an exact exponent identity in all spatial dimensions. In particular, the correlator, $C_0(k) = 2\Gamma_c |G_0(k)|^2$, scales as $k^2$ because of the conservative nature of the noise. It gets a contribution to one-loop order of $k^4$, as one vertex scales like $W \sim k^2$ (Fig. 3). The same applies to the vertex renormalization. More interestingly, the propagator does not get renormalized either, meaning that mean field theory is exact at any dimension [11]. We then have $z = 2$ and substituting this in the renormalisation flow for the the correlator we obtain $\chi = -\frac{d}{2}$. Given that $\langle\rho(x,t)\rho(x',t)\rangle = |x - x'|^{2\chi}F(x/t^z)$, we find that the equal time two-point connected correlation function should scale as $c(x,x') \sim |x-x'|^{-1}$ for $d = 1$.

#### 5.2 Renormalization group analysis of the full non-local kinetics in sequence space

To compute correlation functions and scaling exponents in sequence space we might try to obtain them directly from the field theory. However, such an approach is not feasible and gives exponents which are not in agreement with our numerical simulations (calculations not shown here). The reasons for this are two-fold: First, the field theory is not massless and can be massless only without de-novo DNA methylation. A theory where the mass term is always non zero is not scale invariant and so we will not be able to find critical exponents as observed experimentally and in the simulation. In order to overcome this problem we may try to do a change of variables in the action Eq.(26), and making a Hopf-Cole transformation around a dynamical mean field solution, such that in that reference frame the theory is massless. Even though the theory will be renormalizable, the exponent does not match our stochastic simulations (calculation not shown here). One reason for the failure of such an approach is that we would need to consider perturbations of any order in the field theory and the a one-loop calculation will not be sufficient. The second and main reason of the failure is that, after expanding around a base state integrals of the form (26) cannot be approximated without losing the length scale, $1/\langle\phi\rangle$, associated with the effective cutoff of interactions.

The key insight to calculate the correlation function is that Eq. (26) gives rise to two spatial regimes in sequence space: for short distances interactions are long-range following a power law decay with exponent 1/3 while for distances much larger than $1/\langle\phi\rangle$ interactions decay exponentially and are effectively local. In the following, we will therefore derive the correlation function separately for these two regimes using renormalisation group methods and we will confirm these results with numerical simulations (Fig.S2A)

##### 5.2.1 Short distance regime

As understood in the previous sections all the local non-linearities are irrelevant under renormalization for conservative noise. Importantly, conservative noise is present not only in physical space, but also in sequence space. This is due to the fact that noise can in general be derived from the field theory (26) by identifying terms that are proportional to $\hat{\phi}^2$. In the expansion of the action these terms are both non-conserative, i.e. proportional to $\hat{\phi}^2$ and conservative, i.e. proportional to $\hat{\phi}^2\partial_s^2\phi$. In order to develop a method that is capable to describe the short-distance regime we consider the action (26) and, after taking the semiclassical approximation, we expand it to first order in $\phi$, and $\partial_s$. After this we obtain

$$\partial_t\phi(s,t) = \int_0^s dy\,\phi(y)|s-y|^{-\lambda}e^{-\int_{z=0}^{s-y} dz\,\phi(z)} + \int_0^s dy\,\partial_y\phi(y)|s-y|^{1-\lambda}e^{-\int_{z=0}^{s-y} dz\,\phi(z)}\,, \quad (57)$$

where $s$ is the position in sequence space. For the sake of brevity we omitted the noise terms and integrals of the same form describing interactions with the right nearest bound site. The interaction kernel Eq. (26) has the form $|s-y|^{-\lambda}e^{-\int_{z=0}^{s-y} dz\,\phi(z)}$, with $\lambda = 1/3$ in the case of de-novo DNA methylation. By considering a perturbation $h(s,t)$ around the mean field solution, $\phi_0(t)$, i.e. $\phi(s,t) = \phi_0(t) + h(s,t)$, Eq. (59) can be expressed to first order as

$$\begin{aligned}\partial_t\phi_0(t)+\partial_t h(s,t) = e^{-\phi_0(t)}\int_0^s dy\,\phi_0(t)|s-y|^{-\lambda}\left[1-\int_{z=0}^{s-y} dz\,h(z)\right]+\\ e^{-\phi_0(t)}\int_0^s dy\,h(y)|s-y|^{-\lambda}\left[1-\int_{z=0}^{s-y} dz\,h(z)\right]+\text{h.o.}\,.\end{aligned}\tag{58}$$

The first two terms on the right hand side cancel with the first one on the left hand side, which is the dynamical mean field solution. Taken together, we find

$$\partial_t h(s,t) = e^{-\phi_0(t)}\int_0^s dyh(y)|s-y|^{-\lambda}\left[1-\int_{z=0}^{s-y} dz\,h(z)\right]+\text{h.o.}\,.\tag{59}$$

After a change of variables, $w = z + y$, we obtain

$$\begin{aligned}\partial_t h(s,t) =& e^{-\phi_0(t)}\int_0^s dy\,h(y)|s-y|^{-\lambda}\\ &- e^{-\phi_0(t)}\int_0^s dy\int_y^s dw\,h(y)|s-y|^{-\lambda}h(w-y)+\xi(s,t)\,.\end{aligned}\tag{60}$$

In this expression we recognize a convolution of a fractional integral of a function and the function itself. As discussed in the beginning of this section, the noise, $\xi(s,t)$, for the perturbation $h(s,t)$ has the same form as derived in the previous section. Specifically, it has both conservative and non conservative terms, $\langle\xi(s,t)\xi(s',t')\rangle = \delta(t-t')(2\Gamma_{NC}-2\Gamma_C\partial_s^2)\delta(s-s')$. $\Gamma_C$ and $\Gamma_{NC}$ are the noise strengths for conservative and non conservative noise, respectively. As a side remark, while in the case of only conservative noise the non-local and non-linear term becomes relevant under renormalisation below a critical dimension $d_c = 2(2-\lambda)$. For only non conservative noise the critical dimension is $d_c = 2(3-\lambda)$. The non linear term is identified as a fractional integral,

$$I^\alpha f = \frac{1}{\Gamma(\alpha)}\int (x-y)^{\alpha-1}f(y)\tag{61}$$

where $\Gamma(\alpha)$ is the gamma function. Upon identifying $\alpha - 1 = -\lambda$ in Fourier space the value of this integral scales as $q^{\lambda-1}$.

In order to regularise the theory we introduce an auxiliary process. The lowest order spatial derivative consistent with the symmetries of the theory is $\partial_x^2\phi$. Taken together, taking into account interactions with the right nearest bound site we obtain in Fourier space

$$\partial_t h(q,t) = \left(e^{-\phi_0(t)}q^{\lambda-1} - q^2\right) h(q,t) - q^{\lambda-1}e^{-\phi_0(t)}h(q,t)^2 + \xi(q,t)\,. \tag{62}$$

From the dynamical mean field solution of the first moment, Eq. (29), we know that $e^{-\phi_0(t)} = e^{\left[-t^{\,1/(1-\lambda)}\right]}$. In the frequency domain we then obtain for small timest $\to 0$ or $\omega \to \infty$,

$$i\omega h(q,\omega) = (q^{\lambda-1} - q^2)h(q,\omega) - q^{\lambda-1}h(q,\omega)^2 + \xi(q,\omega)\,. \tag{63}$$

As a side remark, the inverse free propagator is $G_0^{-1} = i\omega + q^2 - q^{\lambda-1}$, which is defined based on the linear part of Eq. (63),

$$\left(i\omega + q^2 - q^{\lambda-1}\right) h(q,\omega) = \xi(q,\omega)\,. \tag{64}$$

and the correlator reads

$$C_0 = \left(2\Gamma_{NC} + 2\Gamma_C q^2\right) |G_0|^2\,. \tag{65}$$

Continuing with our analysis, the general solution of Eq. (63) obtains the form

$$h(q,\omega) = \frac{q^{1-\lambda}}{2J}\left(-q^2 + q^{1-\lambda} + \sqrt{4q^{1-\lambda}\xi + (q^2 - q^{1-\lambda}) + i\omega} - i\omega\right)\,. \tag{66}$$

In Fourier space, two-point correlation functions are defined as $\langle h(q,\omega)h(q',\omega')\rangle$. In the short time and large wavelength limit we keep the lowest order in $q$. Transforming back to real space and setting $t = t'$ we find that

$$\langle h(s,t)h(s',t)\rangle = \left[2\left(|s-s'|^2\Gamma_C + \Gamma_{NC}\lambda(1+\lambda)\right)|s-s'|^{-2-\lambda}\cos(\pi\lambda)/2\Gamma(\lambda)\right] , \quad (67)$$

where $\Gamma(\lambda)$ is the gamma function. We therefore obtain for the scaling of the correlation function $\langle h(s,t)h(s',t)\rangle \sim |s-s'|^{-\lambda}$ with $\lambda = 1/3$ in the case of *de-novo* DNA methylation.

We expect that higher order corrections to this result will depend on the parameters of the model, in particular on average enzyme occupancy, $\phi_0$. In order to understand this point, we may reason that the previous derivation is exact at low values of the average occupancy, $\phi_0$, whilst for larger values of $\phi_0$ we expect higher order corrections to become relevant. As explained above, in the context of spacial correlations we use the terms enzyme occupancy and DNA methylation synonymously.

To understand such higher order corrections it is convenient to temporarily consider the correlation function in non-renormalised physical space, and then transform back to sequence space. We begin by noting that in sequence space the correlation function decays as $|s-s'|^{-1/3}$ for vanishing values of averager form, local DNA methylation. In non-renormalised physical space for vanishing average DNA methylation the correlation function must scale in the same way as in sequence space, i.e. $\sim |\xi - \xi'|^{-1/3}$. We now consider the effect of $n$ DNA methylation events. According to Eq. (40), the this leads to a contraction $\Delta\xi' = \Delta\xi^{-(1/3)^n}$. Going back to sequence space, where length elements scale as $\Delta s \sim \Delta\xi^3$ we obtain that the correlation function decays as $|s-s'|^{-(1/3)^{n+1}}$. $n$ is a monotonically increasing function of the local average DNA methylation level which vanishes for $\phi_0 \to 0$. Expanding to first order we obtain approximately $n \approx \alpha\phi_0 + \dots$ where $\alpha$ is a parameter that we determined to be approximately equal to 1 numerically. Correlation functions then scale as $\langle h(s,t)h(s',t)\rangle \sim |s-s'|^{-(1/3)^{1+\phi_0}}$.

We can already notice from the bare propagator and correlator in Eq (65) that in the short wavelength regime diffusion takes over and in case of conservative noise we expect correlation functions to be described by other exponents. By dimensional argument as in the previous section it is possible to realize that there is another value for the critical exponent, $\chi = -(1+d+\lambda)/3$, which follows from taking into account higher order non linearities and it will be the correct exponent for the long tail of the correlation functions. In the following we are going to prove this simple scaling argument with renormalization group methods.

##### 5.2.2 Long distance regime

In order to calculate the exponents for the long distance regime we begin with Eq. (59) which, as above, is regularised by including the lowest order spatial derivative that is agreement with the model symmetries. After linearisation we obtain

$$\begin{aligned} \partial_t h(s) =& \partial_s^2 h(s) + \int_0^s dy h(y)|s-y|^{-\lambda} + \\ & - \int_0^s dy h(y)|s-y|^{1-\lambda}(h(s) - (s-y)\frac{1}{2}\partial_s h(s)) + \xi(s,t)\,. \end{aligned} \tag{68}$$

where we used $\int_a^b dx\, f(x) \approx (b-a)(f(a)+f(b))/2$.

Before proceeding with renormalisation, we have to take into account other possible non-linearities. We started by considering the linear order in $\phi$ and we obtained an equation(Eq. (68)) involving quadratic terms, $\phi^2$, due to the expansion of the integral. We must therefore come back to the field theory Eq. (26) and keep quadratic terms as well. The only quadratic term in the field theory is

$$\left(1-\hat{\phi}(s)\right)\left(-\phi(s)\hat{\phi(s)}\right)\left[\int_0^s \mathrm{d}y \frac{\hat{\phi}(s-y)\phi(s-y)}{y^\lambda} e^{-\int_{z=0}^{y}\mathrm{d}z\hat{\phi}(s-z)\phi(s-z)}\right]. \tag{69}$$

After functional minimization it with an opposite sign, such that both terms cancel out. This is not surprising, because the symmetry of the system we are studying would not allow a term that breaks the space reversal symmetry $s \to -s$. Taken together, we find

$$\partial_t h(s) = \partial_s^2 h(s) + \int_0^s dy h(y)|s-y|^{-\lambda} + \frac{1}{2}\int_0^s dy h(y)|s-y|^{2-\lambda}\partial_s h(s) + \xi(s,t)\,. \quad (70)$$

Considering both right and left nearest neighbour interactions, the advective terms cancel out. Including the next highest order term we obtain

$$\partial_t h(s) = \partial_s^2 h(s) + \int_0^x dy h(y)|s-y|^{-\lambda} + \frac{1}{2}\partial_s^2 h(s)\int_0^s dy h(y)|s-y|^{2-\lambda} + \xi(s)\,. \quad (71)$$

We can generalize to any spatial dimension by considering the previous equation with a spatial coordinate in vector form, $\mathbf{s}$. In Fourier space the previous equation can then be written in compact form,

$$G_0(\mathbf{q})^{-1} h(\mathbf{q},\omega) = \xi(\mathbf{q},\omega) - \nu\int_{\mathbf{k}} W(\mathbf{q},\mathbf{k})h(\mathbf{q},\omega)h(\mathbf{k}-\mathbf{q},\omega)\,, \quad (72)$$

where $h(\mathbf{q},\omega) = \int d\mathbf{s}\int dt h\,(\mathbf{s},t)\,e^{i\mathbf{q}\mathbf{s}}e^{i\omega t}$, $G_0^{-1} = \left(i\omega + D_0\mathbf{q}^2 + J|\mathbf{q}|^{-\lambda}\right)$. We reintroduced the dimensional parameters from the adimensional Eq. (72) as we are interested in how they scale under renormalization. In Eq. (72) we defined the vertex, which accounts for the non linear part, as

$$W(\mathbf{q},\mathbf{k}) = \frac{\nu}{2}\left[\frac{\mathbf{q}(\mathbf{k}-\mathbf{q})}{|\mathbf{k}-\mathbf{q}|^{3-\lambda}} + \frac{(\mathbf{k}-\mathbf{q})\mathbf{q}}{|\mathbf{q}|^{3-\lambda}}\right]\,. \quad (73)$$

In the limit $\mathbf{k} \to 0$ (hydrodynamic limit) it scales as $\mathbf{k}$, which implies non renormalization of the vertex function. The RG flow then has a general structure of the form

$$\begin{aligned}\partial_l D_0 &= [z-2+A_{D_0}]\, D_0\,,\\ \partial_l \nu &= [z+\chi-2+(3-\lambda)]\,\nu\\ \partial_l \Gamma_C &= [z-2\chi-d-2+A_{\Gamma_C}]\,\Gamma_C\\ \partial_l \Gamma_{NC} &= [z-2\chi-d+A_{\Gamma_{NC}}]\,\Gamma_{NC}.\end{aligned} \tag{74}$$

As we are interested in the short wavelength and so long distances, we set $J=0$ and consider rescaling of the other quantities.

#### 5.3 One Loop correction

After performing standard renormalization group procedures as outlined in the previous section, with the same diagrams as in Fig. 3, we recover the RG flow for the parameters,

$$\begin{aligned}\partial_l D_0 &= \left[z-2-\frac{K_d\nu^2}{dD_0^3}\left[(d-2)\,\Gamma_{NC}+(d-3)\,\Gamma_C\right]\right] D_0\,,\\ \partial_l \nu &= [z+\chi-2+(3-\lambda)]\,\nu\,,\\ \partial_l \Gamma_C &= \left[z-2\chi-d-2-\frac{K_d\nu^2}{2dD_0^3\Gamma_C}(1+d)(\Gamma_{NC}+\Gamma_C)^2\right]\Gamma_C\,,\\ \partial_l \Gamma_{NC} &= [z-2\chi-d]\,\Gamma_{NC}\,,\end{aligned} \tag{75}$$

where $K_d = S_d/(2\pi)^d$ and $S_d$ is the area of a $d$ dimensional sphere.

From the non renormalization of the non conserved noise and of the couplings we get the exact exponent identities $\chi = (-1-d+\lambda)/3$ and $z = (-2+d+2\lambda)/3$. In d=1 and for long distances correlations then decay with an exponent $2\chi = -10/9$. By performing the same procedure as for the short tail, we can estimate the lowest order correction given by finite values of local average DNA methylation. In this case correlation functions for the long tail would scale as $|s-s'|^{-(10/9)^{1+\langle m\rangle}}$.

Taken together, we find that the correlation function in sequence space decays in two algebraic regimes,

$$C(s-s') = \begin{cases} |s-s'|^{-\left(\frac{1}{3}\right)^{1+\langle m\rangle}}, & \text{for } |s-s| \ll 1/\langle m\rangle\,, \\ |s-s'|^{-\left(\frac{10}{9}\right)^{1+\langle m\rangle}}, & \text{for } |s-s| \gg 1/\langle m\rangle\,. \end{cases} \tag{76}$$

The cross-over between these regimes stems from an effective exponential cutoff of the long-range interactions. The position of the cross-over scales with the only length scale in the system, the typical distance between neighbouring methylated CpGs, $1/m$, and, intuitively, separates a regime dominated by active feedback between DNA methylation and topology and a regime characterised by passive, conservative fluctuations. The numerical prefactor of the proportionality between the position of the crossover and $1/\langle m\rangle$ depends on the statistics of distances between neighbouring CpGs in base pairs. We determined this prefactor using stochastic simulations of the disordered system representing the actual CpG positions in the mouse genome (see below) and found that the position of the crossover is approximately equal to $350/\langle m\rangle$.

### 6 Derivation of cross-correlation functions

To derive the cross-correlation between DNA methylation we begin with the Master equation (2) and introduce a complementary binary vector $\mathbf{a}$, $a_i \in \{0,1\}$, which describes whether a site $i$ is accessible ($a_i = 1$) or not ($a_i = 0$). For notational simplicity we assume $i < j$. We then couple this vector to the DNA methylation dynamics in the simplest form compatible with our model interpretation in physical space. We begin by considering the expectation value of the product $m_i a_j$,

$$\langle m_i a_j \rangle = P(m_i = 1, a_j = 1) = P(a_j = 1 | m_i = 1) P(m_i = 1)\,. \tag{77}$$

$P(a_j = 1|m_i = 1)$ cannot be computed directly as it implicitly depends on other values of $\mathbf{a}$ and $\mathbf{m}$. To proceed, we therefore in a first step "integrate in" the random variable describing accessibility at position $a_{j-1}$,

$$\langle m_i a_j \rangle = \sum_{a_{j-1}} P(a_j | a_{j-1}, m_i) P(a_{j-1} | m_i) P(m_i)\,. \tag{78}$$

Applying this step for the second factor we find

$$\langle m_i a_j \rangle = \sum_{a_{j-1}, a_{j-2}} P(a_j = 1 | a_{j-1}, m_i = 1) P(a_{j-1} | a_{j-2}, m_i) P(a_{j-2} | m_i = 1) P(m_i = 1)\,. \tag{79}$$

Repeating these steps $|j - i|$ times, for the sake of simplicity writing $P(m_i = 1)$ as $P(m_i)$), we obtain

$$\langle m_i a_j \rangle = \sum_{a_k, i \leq k < j} P(a_j | a_{j-1}, m_i) P(a_{j-1} | a_{j-2}, m_i) \cdot \ldots \cdot P(a_{i+1} | a_i, m_i) P(a_i | m_i) P(m_i)\,. \tag{80}$$

We are still not in a position to give physical expression for the conditional probabilities as they have an unknown dependence on $m_i$. For simplicity, we for now assume that the mechanical coupling between base pairs is much stronger than the coupling of DNA methylation marks,

$$\langle m_i m_{i+1} \rangle \ll \langle a_i a_{i+1} \rangle\,. \tag{81}$$

Therefore, $P(a_j | a_{j-1}, m_i) \approx P(a_j | a_{j-1})$ for $i \neq j$. We will elaborate on this assumption in more detail below.

In the model, the probability of binding decays according to Eq. (31). The physical interpretation of this kernel was that it reflects the binding probability of enzymes a site being compacted. Therefore, in our model, the probability that a site is compacted decays with the distance to the nearest methylated site to the power of $-1/3$.Therefore, in order to reflect this kernel, the conditional probability $P(a_j|a_{j-1})$ should decay as $P(a_j|a_{j-1}) \propto a_{j-1}(1 - K_a/k^\lambda)$ (where $a_{j-1}$ plays the role of a delta function). Taken together, we then obtain

$$\langle m_i a_j \rangle = P(m_i)P(a_i|m_i) \prod_{k=1}^{|j-i|} \left(1 - \frac{K_a}{k^\lambda}\right) . \tag{82}$$

In our derivation we implicitly assumed that the conditional probabilities $P(a_j|a_{j-1})$ do not explicitly depend on DNA methylation values at positions other than $i$. In principle, in order to obtain simple expressions for the conditional probabilities we would have had to not only sum over intermediary positions in the accessibility vector, $\{a_k\}$, but also over all positions in the DNA methylation vector $\{m_k\}$, ultimately giving a sum over exponentially weighted paths between $m_i$ and $a_j$ (Fig. 4). In the limit, where the mechanical coupling $K_a$ is much stronger than the coupling between DNA methylation events this sum over exponentials is dominated by the path with the highest contribution of $K_a$, the orange path in Fig. 4.

With $P(m_i) = \langle m_i \rangle$ and defining the local coupling between DNA methylation and accessibility $\alpha = P(a_i|m_i)$ we finally obtain

$$\langle m_i a_j \rangle = \alpha \langle m \rangle \exp\left(-K_a |i-j|^{2/3}\right) . \tag{83}$$

In summary, we find that the strength of cross-correlation is linearly proportional to the average DNA methylation level. By contrast, the length scale of the decay, $K_a^{3/2}$, is independent of average DNA methylation. Both results, as well as the functional

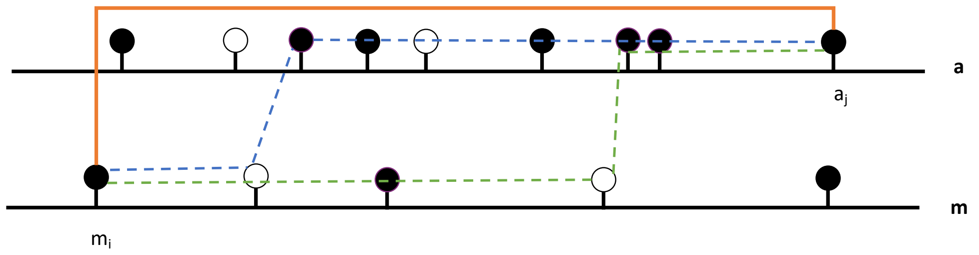

Figure 4: Possible paths from $m_i$ to $a_j$. The orange solid path is the dominant contribution

form of the cross-correlation function, are in good agreement with the experimental data, Fig. S3.

### 7 Stochastic simulations

To test the validity of our analytical results and their dependence on disorder in CpG positions we performed extensive stochastic simulations. These simulations were performed by integration of the master equation Eq.(1) using Gillespie's algorithm [7]. For the reasons outlined above linear and uncorrelated processes do not influence the exponents. Therefore, we considered no unbinding and demethylation processes. We used dimensionless units where the binding and methylation rates are set to $(J, k_m) = (1, 1)$, respectively. We performed simulations on a one dimensional lattice, where for the distribution of distances between neighbouring CpGs was sampled from a distribution resembling the empirical distribution taken from chromosome 1 of the mouse genome. For all simulations we set the lattice size to $10^6$. The simulation of Eq. (51) (Fig. 2F of the main text) are performed with the package xmds2 [13], which performs a pseudospectral integration of the stochastic PDE.

[13] Dennis, Graham R. Hope, Joseph J. Johnsson, Mattias T. , XMDS2: Fast, scalable simulation of coupled stochastic partial differential equations *Computer Physics Communications* **184**

[14] Cross, M. C. and Hohenberg, P. C.,Pattern formation outside of equilibrium *Rev. Mod. Phys.* **65**,
